# Emotional valence modulates the topology of the parent-infant inter-brain network

**DOI:** 10.1101/623355

**Authors:** Lorena Santamaria, Valdas Noreika, Stanimira Georgieva, Kaili Clackson, Sam Wass, Victoria Leong

**Affiliations:** Department of Psychology, University of Cambridge; School of Psychology, University of East London; Division of Psychology, Nanyang Technological University, Singapore

**Author notes:** Corresponding Author: Victoria Leong, Department of Psychology, Downing Street, Cambridge CB2 3EB, U.K.

**Keywords:** EEG hyperscanning, network connectivity, graph theory, emotional expression, mother-infant interaction

## Abstract

Emotional communication between parents and children is crucial during early life, yet little is known about its neural underpinnings. Here, we adopt a dual-brain connectivity approach to assess how emotional valence modulates the parent-infant neural network. Fifteen mothers modelled positive and negative emotions toward pairs of objects during social interaction with their infants (aged 10.3 months) whilst their neural activity was concurrently measured using dual-EEG. Intra-brain and inter-brain network connectivity in the 6-9 Hz (infant Alpha) range was computed during maternal expression of positive and negative emotions using directed (partial directed coherence) and non-directed (phase-locking value) connectivity metrics. Graph theoretical metrics were used to quantify differences in network topology as a function of emotional valence. Inter-brain network indices (Density, Strength and Divisibility) consistently revealed that the integration of parents’ and childrens’ neural processes was significantly stronger during maternal demonstrations of positive than negative emotions. Further, directed inter-brain metrics indicated that mother-to-infant directional influences were stronger during the expression of positive than negative emotions. These results suggest that the parent-infant inter-brain network is modulated by the emotional quality and tone of dyadic social interactions, and that inter-brain graph metrics may be successfully applied to examine these changes in interpersonal network topology.

## INTRODUCTION

### 1.1 Intra-individual neural networks for emotional processing

Emotional processing and regulation involve both ‘top-down’ and ‘bottom-up’ processes of control and regulatory feedback that engage the fronto-limbic network (FLN) (Ochsner et al., 2009). Within the FLN, it is the dorsolateral, ventrolateral and medial prefrontal cortices along with limbic structures, such as the amygdala and hippocampus that have been most commonly implicated in emotion processing and regulation (Ochsner et al., 2009). The basal ganglia are also implicated in the processing of facial (Adolphs, 2002) and vocal (Kotz et al., 2003) emotional expressions. For example, deep brain stimulation of the basal ganglia causes impairment of emotion perception from facial and vocal expressions (Péron et al., 2010). More recent approaches have used connectivity-based measures to examine how these networks of brain regions coordinate their activity dynamically during emotion processing (Diano et al., 2017; Sato et al., 2017). Extensive previous research has also examined how these intra-individual neural networks become disrupted during atypical development (Goulden et al., 2012; Lu et al., 2012; Nicholson et al., 2017).

Neural oscillations (which are measurable using scalp EEG) reflect rhythmic fluctuations in the synchronization of neuronal populations at a millisecond timescale. Activity in the EEG Alpha band is strongly implicated in the processing of emotional stimuli and social cognition in adults and infants (Allen et al., 2018; Coan & Allen, 2004). Studies with normal adults indicate that Alpha power over the left and right frontal brain regions respond differentially to emotional valence (Davidson, 1984, 1998). Activation over the left frontal area is commonly associated with the experience of positive emotions such as joy or interest, whereas right frontal Alpha EEG power is associated with disgust, crying and sadness. Further, individuals who experience mood disorders such as depression exhibit atypical patterns of EEG asymmetry, commonly showing higher right frontal EEG activity than controls (Gotlib et al., 1998). Recent research has also started to examine intra-individual network topology during emotion processing using graph-theoretic measures. For example, a recent study with adults showed that EEG graph-theoretic features performed better than traditionally used EEG features (such as spectral power and asymmetry) on the automatic classification of affective neural states (Gupta et al., 2016).

Behavioral and neuroimaging studies into early development suggest that the neural architecture for the detection and prioritized processing of emotional expressions, such as fear, emerges sometime during the first year of life (Hoehl, 2013; Hoehl et al., 2019; Leppänen & Nelson, 2009). It is during this time that infants’ visual system is sufficiently developed to support the discrimination of most facial expressions (Leppänen & Nelson, 2006), and that infants begin to exhibit a reliable attentional bias towards fearful facial expressions (Nelson & De Haan, 1996). For example, 7-month-old infants look longer at fearful than happy facial expressions (Nelson et al., 1979) and are slower to disengage their attention from a fearful face than happy or neutral faces (Leppänen et al., 2010). Recordings using EEG responses to facial expressions in 7-month-old infants have shown that particular ERP components are enhanced when infants view fearful facial expressions (Hoehl & Striano, 2008; Leppänen et al., 2007). Influences of the early environment on infants’ neural processing of emotion have also been shown: whereas typically-developing infants exhibit greater left versus right frontal brain activity, infants of depressed mothers exhibit greater right frontal brain asymmetry (Dawson et al., 1992; Field et al., 1995; Field et al., 1995). Understanding the early development of emotion processing is considered essential, in particular, for helping to identify and intervene in cases of atypical development.

### 1.2 Inter-individual neural networks for emotional processing

Research is increasingly moving from approaches that emphasise localised structure-function correspondences towards a more distributed, network-based approach that studies how activity is coordinated across multiple brain regions on an intra-individual basis (Bullmore & Sporns, 2009). Less well established, however, is research into how network-based patterns of brain activity subsist *between* individuals during human interactions – i.e. on an *inter*-individual basis (Schilbach et al., 2013).

Inter-individual dynamics play an essential role in many forms of human interaction – particularly so during early development (Feldman, 2007; Jaffe et al., 2001). Social co-ordination between parents and their offspring engenders early learning across multiple domains of social and cognitive development (Csibra & Gergely, 1998; Feldman, 2007; Rogoff, 1990). The behaviour of human infants and their adult caregivers is closely co-ordinated, and adult-infant temporal contingencies occur across behavioural, physiological and neural domains. For example, adults’ and children’s gaze patterns (Kaye & Fogel, 1980), vocalisations (Jaffe et al., 2001), emotional states (Cohn & Tronick, 1988), autonomic arousal (Skoranski et al., 2017), hormonal fluctuation (Spangler, 1991), and neural oscillatory activity (Leong et al., 2017) all show mutual temporal dependencies of different forms.

Dyadic neuroimaging studies with adults using functional Near-Infrared Spectroscopy (fNIRS) have shown that during verbal (non-emotional) communication, dyads develop synchronous patterns of activity between brain regions such as the inferior frontal gyrus, prefrontal and parietal cortices (Jiang et al., 2012). A recent fNIRS study examining parents and their 7.5-year-old children found that their prefrontal regions showed synchronisation during conditions of social co-operation (but not during competition), and that the degree of neural synchronisation mediated the relationship between parent’s and children’s emotional regulation abilities, as assessed via questionnaires (Reindl et al., 2018). Using EEG to examine synchronisation at a finer time-scale, we have also shown that neural synchronicity occurs between adults and infants in the Theta and Alpha bands, and that such dyadic neural connectivity is modulated by eye contact between the adult speaker and infant listener (Leong et al., 2017). Other research has, similarly, used EEG to explore inter-personal dynamics between adults (Babiloni & Astolfi, 2014; Dumas et al., 2011), including research that has examined larger sized groups (Dikker et al., 2017).

### 1.3 Graph connectivity in two-person neuroscience

The interpersonal neural network contains crucial information with regard to teamwork/co-ordination (Babiloni et al., 2011; Dikker et al., 2017), communicative efficacy (Hasson et al., 2012; Jiang et al., 2012), and social status (e.g. leader-follower relationships; Jiang et al., 2015; Sänger et al, 2012, 2013), but it is not clear exactly how such information is encoded within social networks. Whereas previous dyadic studies have typically focussed on how *strongly* connected the interpersonal network is (i.e. how much information is shared between partners), relatively little previous research has examined the organisation and *topology* of the network itself (i.e. how information flows between partners). This distinction is important because socioemotional factors may modulate the structure of a neural network without necessarily changing its mean strength or activation level. For example, Betzel et al (2017) showed that individual variations in mood and surprise were correlated with changes in neural network *flexibility* (that is, the reconfiguration of network community structure over time). Positive moods were associated with higher levels of network flexibility whereas increased levels of surprise were associated with lower network flexibility.

Graph connectivity measures are useful for capturing the topological properties of neural networks (Bullmore & Sporns, 2009). However, graph metrics are not commonly adapted for use with *inter*-personal neural networks (although see Astolfi et al., 2010, 2015). It has recently been suggested that these network metrics may in fact be usefully applied to multilayer network models in order to understand how information is shared between individuals and across social networks (Falk & Bassett, 2017).

### 1.4 Study overview

Here, we use graph theoretical indices to assess the topology of parents’ and infants’ *intra*-and *inter*-brain neural networks during emotional processing. To study emotional processing, we used a classic social referencing task (Walden & Ogan, 1988) that involved maternal demonstrations of positively-and negatively-valenced emotion. During social referencing, the partner’s social interpretation of events is used to form one’s own understanding of a situation (Feinman, 1982). Social referencing develops over the first year of life, and by 10-12 months of age, infants will seek information from others in novel situations and will use this information to regulate their own affect and behaviour (Feinman et al, 1992). For example, infants at this age will avoid crossing a short visual cliff (Sorce et al., 1985), show less interaction with toys (Gunnar & Stone, 1983; Hornik et al., 1987) and be less friendly to strangers when their mothers model negative emotions as compared to neutral or happy emotions (Feinman & Lewis, 1983; Feinman & Roberts, 1986).

Research has shown that, even in young infants, the brain responds differentially to objects as a function of how other people are reacting to them (Hoehl et al., 2008). Infants’ neural processing of novel objects is enhanced by a fearful, but not a positive, face gazing toward the object (Hoehl & Striano, 2010) – influence that may be enhanced or reduced by infants’ temperamental predisposition (Aktar et al., 2016). However, little is known about the dynamic, *inter*-personal neural mechanisms that support social referencing and emotional co-ordination between parents and their children.

As we were particularly interested in the *direction* of information flow between parents and their children, we assessed network connectivity using both directed (partial directed coherence, PDC) and non-directed (phase-locking value, PLV) indices. We were primarily interested in whether, and how, the topology of *inter*-brain and *intra*-brain networks would be influenced by emotional valence.

## 1 METHODS

### 2.1 Participants

Fifteen mother-infant dyads participated in the study (8 male, 7 female infants). Infants were aged 315.6 days on average (SEM = 9.42 days). All mothers reported no neurological problems and normal hearing and vision for themselves and their infants. The study was approved by the Cambridge Psychology Research Ethics Committee and parents provided written informed consent for them and on behalf of their infants.

### 2.2 Materials

Four pairs of ambiguous novel objects were used. Within each pair, objects were matched to be globally similar in size and texture, but different in shape and colour. Ambiguous novel objects were chosen to ensure that infants would not have previous experience with these objects.

### 2.3 Task protocol

A classic social referencing task was used, which involved positive and negative maternal emotional demonstrations toward novel toy objects (Hirshberg & Svejda, 1990; Hornik et al., 1987; Walden & Ogan, 1988). Infants were seated in a high chair, and a table was positioned immediately in front of them (see Figure 1). Parents were seated on the opposite side of the table, directly facing the infant. The width of the table was 65cm. Each experimental trial comprised a maternal demonstration phase involving one pair of novel objects, and a response phase. Trials began when the mother attracted her infant’s attention by saying “Look”, or by holding one of the objects up. During the demonstration phase, mothers were instructed to show positive affect toward one object and negative affect toward the other object, as illustrated in Figure 1. Mothers were instructed to limit their speech to simple formulaic verbal statements per object (which they repeated for each object), and to model positive or negative emotions in a prescribed manner (e.g. smiling versus frowning) (see Figure 1). The order of object presentation (positive or negative) was counterbalanced across trials, and the order of objects was counterbalanced across participants. Up to 16 trials were presented to each infant and on average, infants completed 9.5 trials (std: 3.6).

**Figure 1.**
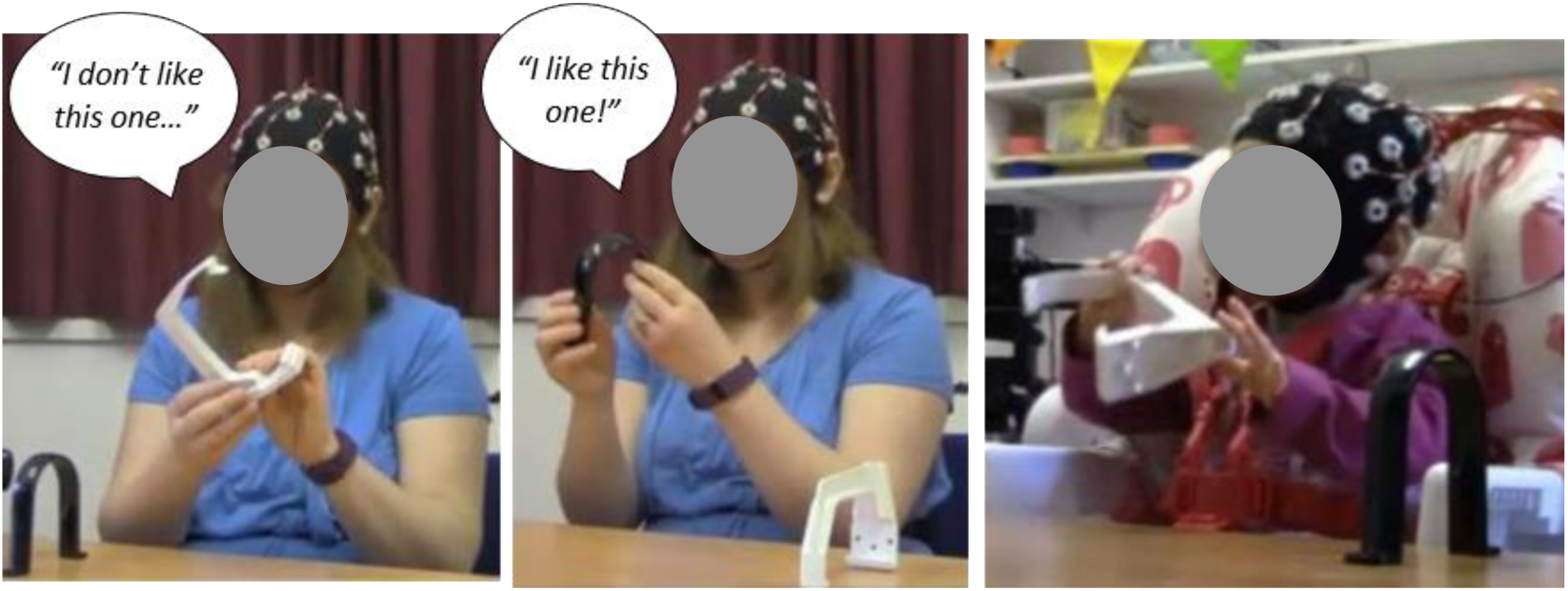
Illustration of experimental setup and task. (Left) Negative object demonstration by adult; (middle) Positive object demonstration by adult; (right) Infant’s interaction with objects. Written informed consent was obtained for the publication of this image.

The period of positive emotion modelling will be referred to as the “*Pos*” condition, and the period of negative emotion modelling will be referred to as the “*Neg*” condition. Across participants, the mean duration of the *Pos* condition was 2.75 seconds (std: 1.26) and the mean duration of the *Neg* condition was 2.48 seconds (std: 1.14). There was no significant difference in the duration between conditions (*p*=0.40, Hedges’g=0.10). Further, as detailed in the Supplementary Materials (S3), there was no significant difference in the mean pitch of maternal utterances between conditions (*p*=0.10, Hedges’g=0.47). However, there was a significant difference in loudness (*p*=0.001, Hedges’g=1.61), where maternal utterances during the *Pos* condition were louder than during the *Neg* condition. In the Supplementary Materials (S3) we provide further analyses controlling for the effect of these acoustic differences on our neural connectivity analyses.

After observing the maternal emotional demonstrations, infants were allowed to interact briefly with the objects before they were retrieved. An experimenter was present throughout the session, but positioned out of the line of sight of both participants, to ensure that participants were interacting as instructed. The experimenter provided new pairs of objects as required, but explicitly avoided making prolonged social contact with either participant.

### 2.4 Baseline task

Each mother-infant dyad also performed a baseline task that did not involve emotional modelling or social interaction. During this baseline task, they were seated in the same configuration as for the main task (across a table from each other), but with a 40 cm high screen in place, so that the infant and adult could see one another, but not the object with which the other was interacting. Mother and infant played with their own toy objects (which were different from the main task). The baseline task was completed either before or after the main task, in a counterbalanced order across participants.

### 2.5 Video recordings

To record the actions of the participants (e.g. start and end of teaching periods), two Logitech High Definition Professional Web-cameras (30 frames per second) were used, directed at the adult and infant respectively. Afterwards, each video recording was manually coded to identify the periods of interest, based on the onset and offset of maternal utterances.

### 2.6 EEG acquisition and pre-processing

#### EEG acquisition

A 32-channel BIOPAC Mobita mobile amplifier was used with an Easycap electrode system for both infant and adult. Electrodes were placed according to the 10-10 international system for electrode placement. Data were acquired using AcqKnowledge 5.0 software, at a 500 Hz sampling rate. The ground electrode was affixed to the back of the neck as this location is the least invasive for infants. The amplifiers for both participants were synchronized through a push button trigger signal that was sent simultaneously to both EEG systems and also simultaneously delivered a LED signal that was visible on both video recordings (for video-EEG synchronisation).

We selected a subset of 16 frontal, central and parietal channels for further analysis (see Figure 2). This sub-selection was done to reduce the computational cost of the analysis, and also because previous research has shown that the contribution of speech myogenic artifacts is relatively stronger at peripheral electrodes (Porcaro et al., 2015; Brooker & Donald, 1980). The selected channels were: F_3_, F_z_, F_4_, FC_1_, FC_2_, C_3_, C_z_, C_4_, CP_5_, CP_1_, CP_2_, CP_6_, P_3_, P_z_, P_4_, and PO_z_.

**Figure 2.**
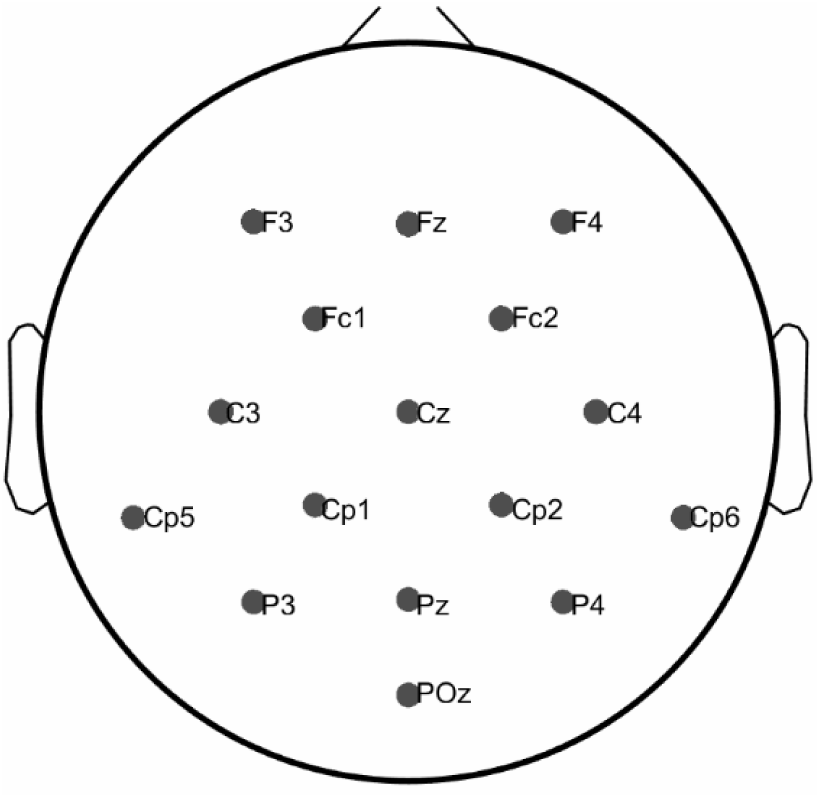
Electrode map of selected channels.

#### EEG pre-processing for motion-related artifacts

EEG signals were band-pass filtered in the range of 1 to 16 Hz in order to suppress line noise as well as minimise as far as possible the effect contamination by muscular (e.g. speech and facial) artifacts which are most prominent at frequencies over 20 Hz (Whitham et al., 2007). Next, a threshold criterion (± 80μV) was applied to remove high-amplitude artifacts. Finally, visual inspection of the data was performed to eliminate residual artifacts. Only EEG segments that were artifact-free across all electrodes for *both* mother and infant within each dyad were used for further analysis (on average 83.62% of the data were used for further analysis).

#### Baselining

Prior to conducting connectivity analysis, and in order to reduce differences in amplitude across participants in the dataset, and between infants and parents, two normalisation steps were applied. First, the main task data from each participant were z-normalised according to their corresponding baseline task data. That is, the mean of the baseline task data was subtracted from the main task data, and the result was then divided by the standard deviation of the baseline task data. Second, the main task data from each dyad were z-normalised relative to one another.

### 2.7 Connectivity metrics (6-9 Hz, infant Alpha band)

Two sets of connectivity calculations were performed. First, we examined intra-brain connectivity, between the individual electrodes in the infant and the adult recordings considered separately. Second, we examined inter-brain connectivity, between the infant electrodes and the adult electrodes. Our analyses focused on assessing network connectivity in the infant Alpha frequency band (6-9 Hz; Marshall et al., 2002) for three reasons. First, because Alpha activity is strongly implicated in the processing of emotional stimuli and social cognition in adults and infants (Allen et al., 2018; Coan & Allen, 2004). Second, because we had previously observed adult-infant neural synchronicity in this frequency range (Leong et al., 2017). Third, because our previous research and that of others has shown that this frequency range is least affected by facial myogenic artifacts (Georgieva et al., under review; Goncharova et al., 2003) and global Alpha network characteristics can be reliably assessed even in 10 month infants (Velde et al., 2019)

We contrasted two measures of neural connectivity: one non-directed measure (Phase Locking Value, PLV) and one directed measure (Partial Directed Coherence, PDC). Both connectivity measures were computed on 6-9 Hz band pass filtered data, using 2s sliding windows with a 50% overlap. All computations were performed using in-house adaptations of functions from publicly available Matlab^©^ based toolboxes (He et al., 2011; Niso et al., 2013).

<u>Phase Locking Value</u> (PLV) measures frequency-specific transients of phase locking independent of amplitude (Lachaux et al., 1999). The instantaneous phase of the signal was calculated using the Hilbert transform. Two signals x(t) and y(t) with instantaneous phases *φ*_*x*_(*t*) and *φ*_*y*_(*t*) are considered phase synchronised if their instantaneous phase difference is constant:

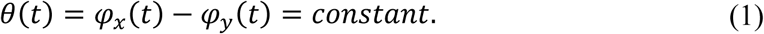

To calculate phase synchronisation, we used PLV defined as:

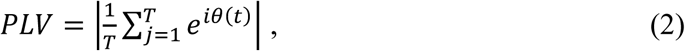

where T is the number of time samples. PLV is a value within the range [0, 1], where values close to 0 indicate random signals with unsynchronised phases and higher values indicate stronger synchronization between the two signals (here, pairs of electrodes).

<u>Partial Directed Coherence</u> (PDC) is based on the concept of Granger Causality (Granger, 1969). It is a spectral estimator and provides the directed influences between each pair of signals in a multivariate data set (Baccala & Sameshima, 2001). If a multivariate data set is understood as an ensemble of simultaneously recorded signals (channels), for a *k*-channel set the model is defined by:

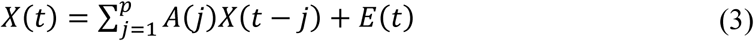

where E(t) is a vector of *k* white noise values for each time point *t. A* is a square *k* × *k* matrix representing the model parameters and *p* is the model order. Transforming the given multivariate model into frequency domain we obtain:

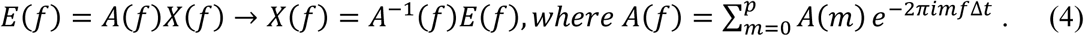

From the transformed model coefficients, A(f), the PDC can be calculated as:

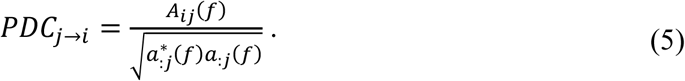

PDC is a normalised measure that can distinguish between direct and indirect connectivity flows better than other Granger causality based metrics such as Direct Transfer Function or its versions (Astolfi et al., 2007).

The application of multivariate models for connectivity analysis requires the estimation of the model order *p*. In this study we implement the Schwarz Bayesian information criterion (SBIC) (Schwarz, 1978) and the Akaike information criterion (AIC) (Akaike, 1974), where the value of the model order was selected based on the measure providing the lowest values across both methods. Under this criterion a model order of 5 was used, which explained the highest proportion of data (91.39% of infants’ data and 81.30% of adults’). To ensure that the implemented model was able to capture the essential dynamics of the data we applied two different techniques to validate the fitted model. First, we calculated the percentage of consistency of the model using the Ding method (Ding et al., 2000). This test provides the percentage of the correlation structure in the data that is captured by the fitted model. 100% of the dataset achieved a consistency of ⇒80%, which is considered to be the acceptable lower limit. Second, the coefficient of determination or r-squared was calculated. This test indicates the percentage of the data that is explained by the model. Again, the entire dataset obtained an *r*-squared value of over 30%, indicating good model estimation (Seth, 2010). The same procedures were used to calculate inter-subject connectivity, where one autoregressive model was created based on the EEG data from the infant-mother dyad.

### 2.8 Statistical validation of connectivity results

#### Intra-brain connectivity

To assess whether the intra-brain connectivity values were significantly above chance, a surrogate data analysis was performed which controlled for spurious (random) connections. To achieve this, a Fourier transform was applied to each data epoch for each channel, and a random permutation of phase values was performed in the frequency domain. Finally, an inverse Fourier transform was used to recreate the surrogate data in the time domain. This process retained the original spectral profile of the data whilst selectively disrupting phase relationships across channels, thereby removing genuine phase-based connectivity patterns. A total of 100 surrogate datasets were created for each participant, channel and epoch. To perform the validation procedure, the neural connectivity indices obtained for the real data were compared against those for the surrogate data at a significance level of *p*<0.05 using paired-samples t-tests, corrected for multiple comparisons using Tukey’s honestly significant difference criterion (Matlab^©^). Individual connections that were not significantly different from their respective surrogates were set to zero (and disregarded for subsequent statistical analyses of differences between conditions).

#### Inter-brain connectivity

To assess whether the measured *inter*-brain connectivity results were significantly above chance, two validation steps were performed. First the analysis using phase-randomised surrogates was performed in an identical fashion to that described above. Second, the neural connectivity values obtained from the real data were compared to a pair-randomised dataset generated by randomly pairing mothers and infants from different sessions whose brain data was non-matching (210 new couples). For this pair-randomised data, any connectivity that existed between the random pairings would have occurred purely by chance (e.g. due to participants experiencing similar environmental conditions during the experiment). As the duration of trials varied across participants, the longer dataset was cropped to the same length as the shorter dataset for each random pair (Reindl et al., 2018). For each condition and EEG channel, a two-sample t-test (significance level of 5% corrected by false discovery rate for multiple comparisons using the Tukey’s honestly significant difference criterion) was performed between the real dataset and the pair-randomised dataset. Individual connections that did not reach significance after the first (surrogate) data step were not included in this second validation step.

### 2.9 Thresholding

Thresholding is necessary to remove spurious connections and to obtain sparsely connected networks, which is a pre-requisite for the computation of many graph metrics (Deuker et al., 2009). Different approaches are used to select an appropriate threshold value. Thresholds can be selected based on the statistics of the data distribution or by taking into account the sparsity of the resulting matrix (Philips et al., 2017). Here, we adopt the most widely used method where a proportional threshold is imposed on all the links within the network. This means that the *density* of the adjacency matrix, defined as the percentage of existing connections with respect to all possible connections in the network, is fixed. Proportional thresholding is expected to lead to more stable networks metrics (Garrison et al., 2015) and is the most widely used technique for studies that compare between experimental conditions or groups (Nichols et al., 2017; Toppi et al., 2012).

To determine the appropriate threshold, we first conducted a visual inspection of the connectivity patterns resulting from thresholding at a range of values (0.17, 0.15, 0.10, 0.08 and 0.05). Figure 3 shows the results that were obtained when different thresholds were applied to the adult grand averaged PLV dataset. By visual inspection of Figure 3, the strongest connections appear to be concentrated within fronto-central scalp regions. As thresholds are relaxed (e.g. above 15%), this pattern becomes increasingly obscured as weaker connections from other regions begin to be included. The same threshold criterion of 15% was applied to the PLV metric as to the PDC data. These selected thresholds offered the most optimal balance between data retention (increased with lower threshold values), readability of connectivity patterns (optimal for higher values) and computational cost (Filho et al., 2016). Section S1 in the Supplementary Materials shows the effect of applying different thresholds for both infant and adult data, and for PDC and PLV metrics, which yielded similar effects to the data shown here.

**Figure 3.**
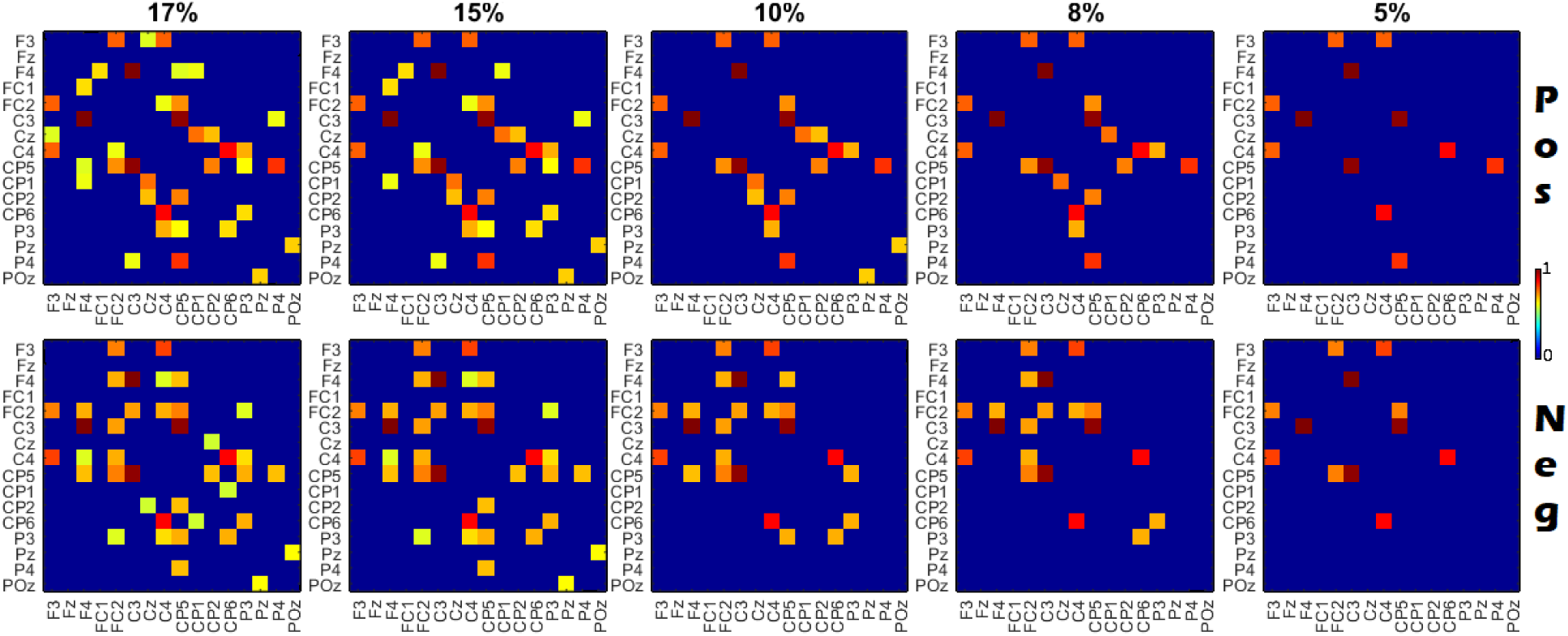
Effect of applying different thresholds to maternal grand average 6-9 Hz PLV matrices. The first row shows the Pos condition and the second row shows the Neg condition. From left to right the threshold values for each column are: 17, 15, 13, 8 and 5% of the strongest links preserved. The threshold of 15% was selected as being the most optimal.

### 2.10 Graph theoretical indices of network topology

A graph consists of a series of nodes (EEG electrodes) and a set of edges (connections) showing the relationships between the nodes. To define a graph, it is necessary to construct an adjacency matrix *A* which captures the connectivity structure of the graph. An adjacency matrix is constructed by comparing the link between each pair of nodes in the connectivity matrix against a corresponding threshold. Edges whose values are larger than the threshold (here, the top 15% as described in the previous section) remained in the adjacency matrix, whilst those with values under the limit were set to 0.

#### 2.10.1 Intra-brain metrics

The indices that define the topology of a network can be broadly divided into four groups: individual metrics (degree, density, strength), functional segregation metrics (clustering coefficient, transitivity, modularity or local efficiency), functional integration metrics (global efficiency, characteristic path length, radius and diameter), and centrality metrics such as betweenness centrality (Rubinov & Sporns, 2010). Here, we selected one metric from each group to provide an overview of these different network properties. As we used a fixed network density (15%), the density and the average degree of the networks would be the same for each experimental condition, hence these indices were not used. Rather, the following indices are reported here:

##### Individual metrics

<u>Strength (S)</u> is the weighted variant of degree. This is typically defined as the sum of neighbouring link weights. In this case, we report the highest value of neighbouring link weights. This is likely to be more informative than mean strength as network density was fixed to retain only the most strongly connected links.

##### Functional segregation

<u>Transitivity (T)</u> is the overall probability for the network to contain interconnected adjacent nodes, revealing the existence of tightly connected communities. In simple terms of network topology, this index represents the mean probability that two vertices that are network neighbours of the same other vertex will themselves be neighbours (Newman, 2003).

##### Functional integration

<u>Global Efficiency (GE)</u> is inversely related to the topological distance between nodes and is typically interpreted as a measure of the capacity for parallel transfer and integrated processing. It is based on the inverse of the shortest path length, which is an indicator of the ease with which each node can reach other nodes within the network using a path that is composed of only a few edges. Hence, the global efficiency is an indicator of the degree to which a network can share information between distributed regions (Kabbara et al., 2018)

##### Centrality

<u>Betweenness centrality (BC)</u> is a measure of centrality. These measures identify central nodes that connect various brain regions. The betweenness centrality of a node is defined as the fraction of all shortest paths in the network that pass through the given node. Nodes with a larger betweenness centrality value will participate in a higher number of shortest paths.

Graph analysis was performed separately for PLV and PDC measures, and for each condition using the Brain Connectivity Toolbox (Rubinov & Sporns, 2010). The resulting intra-brain graph indices were assessed separately for parents’ and infants’ data using Repeated Measures ANOVAs, taking Condition (*Pos* and *Neg* teaching) as a within-subjects factor. Results were corrected for multiple comparisons using Tukey’s honestly significant difference criterion (Matlab^©^).

#### 2.10.2 Inter-brain metrics

Due to the difference in format between individual and inter-brain adjacency matrices, an adaptation process was needed before inter-brain graph indices could be computed. Each dual-brain adjacency matrix comprises of four different sections; the first section of the first N rows and N first columns describes the intra-brain connections for the first member of the dyad (here, the mother). The first N rows and last N columns represent the connectivity *between* the first and second member of the dyad (mother to infant). The last N rows and the last N columns represent the intra-brain connectivity values for the second member (infant) and the last N rows and first N columns represent connectivity between the second and first member of the dyad (infant to mother). For non-directional indices (e.g. PLV), mother-to-infant and infant-to-mother connectivity patterns are symmetric.

Two inter-brain-adapted graph metrics were used here: Strength and Divisibility. These graph metrics were computed for each dyad and experimental condition using only significant inter-brain connections. To maintain an equal density across experimental conditions, the least number of significant connections across both conditions was used in the inter-brain graph analysis – this was 10 connections.

<u>Strength:</u> is the sum of neighbouring link weights as described previously. The adapted version for inter-brain connectivity was calculated as follows:

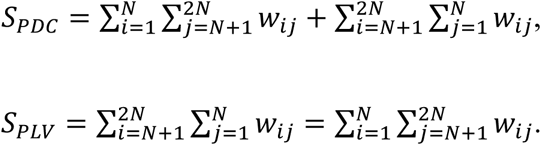

where N is the number of channels for each subject and *w*_*ij*_ is the weight of the connection between node *i* of subject 1 and node *j* of subject 2. As PLV is a non-directed metric, the connectivity matrix from subject 1 to subject 2 is identical to the matrix from subject 2 to 1. For PDC however, separate calculations were performed to assess the <u>directed strength</u> from mothers to infants (*MtoI*) and vice versa, from infants to mothers (*ItoM*):

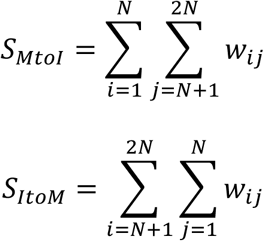

<u>Divisibility:</u> is a measure of how well the entire connectivity network (including intra-and inter-brain connections) can be divided into two sets of nodes, corresponding to the brain of each member of the dyad (Astolfi et al, 2015; 2010; De Vico et al, 2010). It is defined as:

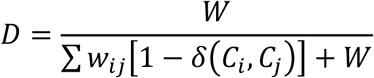

where W is the total weight of the network (including within and inter-brain subnetworks), Ci and Cj indicate the community (which brain) the nodes *i* and *j* belong to respectively. The function δ is binary with values 0 or 1 (1 if vertices *i* and *j* are in the same community and 0 otherwise). The resulting values of D (divisibility) range between [0,1]. For example, in a fully connected network (where all possible links are connected with a value of 1), the resulting value D is 0.67. When the network is fully disconnected (all possible links are set to 0), the resulting D value is 0. A value of D=0.5 is obtained when all inter-brain connections are fully connected (=1), but all within brain connections are disconnected (=0), since in this case the index reduces to D=W/(W+W)=0.5. Conversely, a value of D=1 is obtained when all inter-brain connections are disconnected (=0), but all within-brain connections are fully connected (=1), in which case D=W/(0+W)=1. Therefore, if 0.5 < D < 0.67, it may be inferred that interbrain connections are *stronger than* within brain connections. For values of 0.67 < D < 1, interbrain connections are *weaker than* within brain connections.

<u>Directed Divisibility</u> For PDC, similarly to the directed strength, separate calculations were performed for incoming and out-going directions of connectivity. This would highlight which partner was “leading” the neural integration process during each condition under study. In order to calculate directed divisibility, one of the inter-brain matrices was set to zero each time. For instance, to calculate the directed divisibility from mothers to infants the quadrant corresponding to connections from infants to mothers was set to zero. Therefore, the total weight *W* from the previous equation was transformed to:

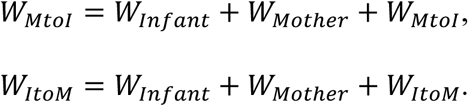

Resulting in the following directed divisibility equations:

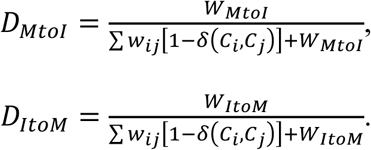

The resulting Strength and Divisibility inter-brain graph indices were subjected to non-parametric Kruskal-Wallis tests (significance level of 5%, multiple comparison corrected using Tukey’s honestly significant difference criterion) to assess statistically significant differences in inter-brain connectivity between conditions (*Pos* and *Neg*).

### 2.11 Intra-and inter-brain density

In addition to the graph metrics of network topology, we also computed measures of intra-and inter-brain network density.

<u>Intra-brain density</u>. Intra-brain density was calculated as the ratio of existing (significant) edges to the total number of possible connections. This index was computed using the non-thresholded data (since thresholds impose a fixed ratio).

<u>Inter-brain density</u>: Here, we defined inter-brain density as an extension of the established within-brain density metric: the ratio of existing (significant) inter-brain edges to the total number of possible interbrain connections. The inter-brain density metric is therefore a measure of neural integration between parents and infants. Calculations were computed over the statistically validated inter-brain connectivity matrices (i.e. to identify significant connections) without any further thresholding. For computation of PLV-based inter-brain density, only one of the inter-brain connectivity matrices was used as mother-to-infant and infant-to-mother matrices are identical. PLV inter-brain density was computed as:

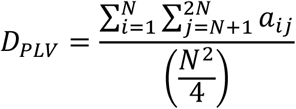

Where *N* is the total number of channels, and *a* represents the existence (or not) of a link between two nodes in the adjacency matrix.

For the PDC measure, both inter-brain connectivity matrices were included and total inter-brain density was computed as the sum of the individual directed densities:

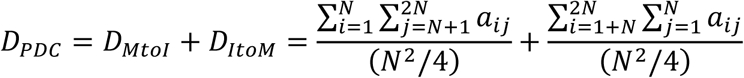

## 2 RESULTS

### 3.1 Intra-brain connectivity

Effective connectivity networks were estimated at the single subject level for each condition (*Pos* and *Neg*) using both non-directed (PLV) and directed (PDC) connectivity metrics.

#### 3.1.1 Intra-brain connectivity by experimental condition

Figures 4 and 5 depict adults’ and infants’ respective grand average connectivity patterns for the 6-9 Hz Alpha band in *Pos* and *Neg* conditions, obtained using PLV (top row) and PDC (bottom row) measures respectively. Stronger connectivity between electrode pairs is indicated with thicker and darker lines.

**Figure 4.**
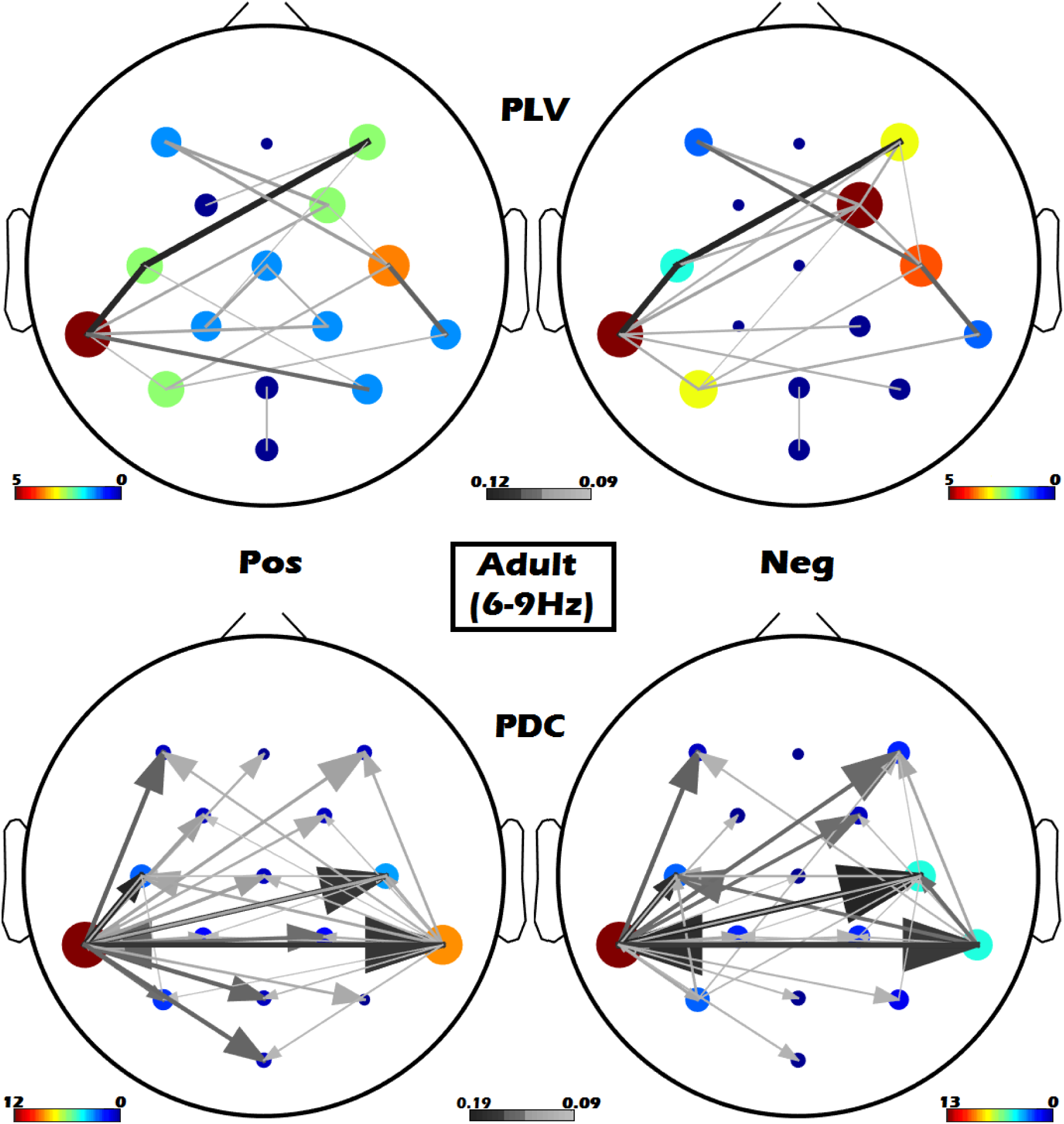
Adult intra-brain connectivity patterns for the 6-9 Hz band using PLV (top row) and PDC (bottom row). The left column shows the Pos condition and right column shows the Neg condition. For each subplot, the colour and size of each node is proportional its degree, where hotter colours indicate higher values and cooler colours indicate lower values. The weight of the edges in the networks are represented in grey scale, where darker colours indicate stronger connections. For the PDC measure, arrows represent the directionality of connections, ending in the node receiving the information flow.

**Figure 5.**
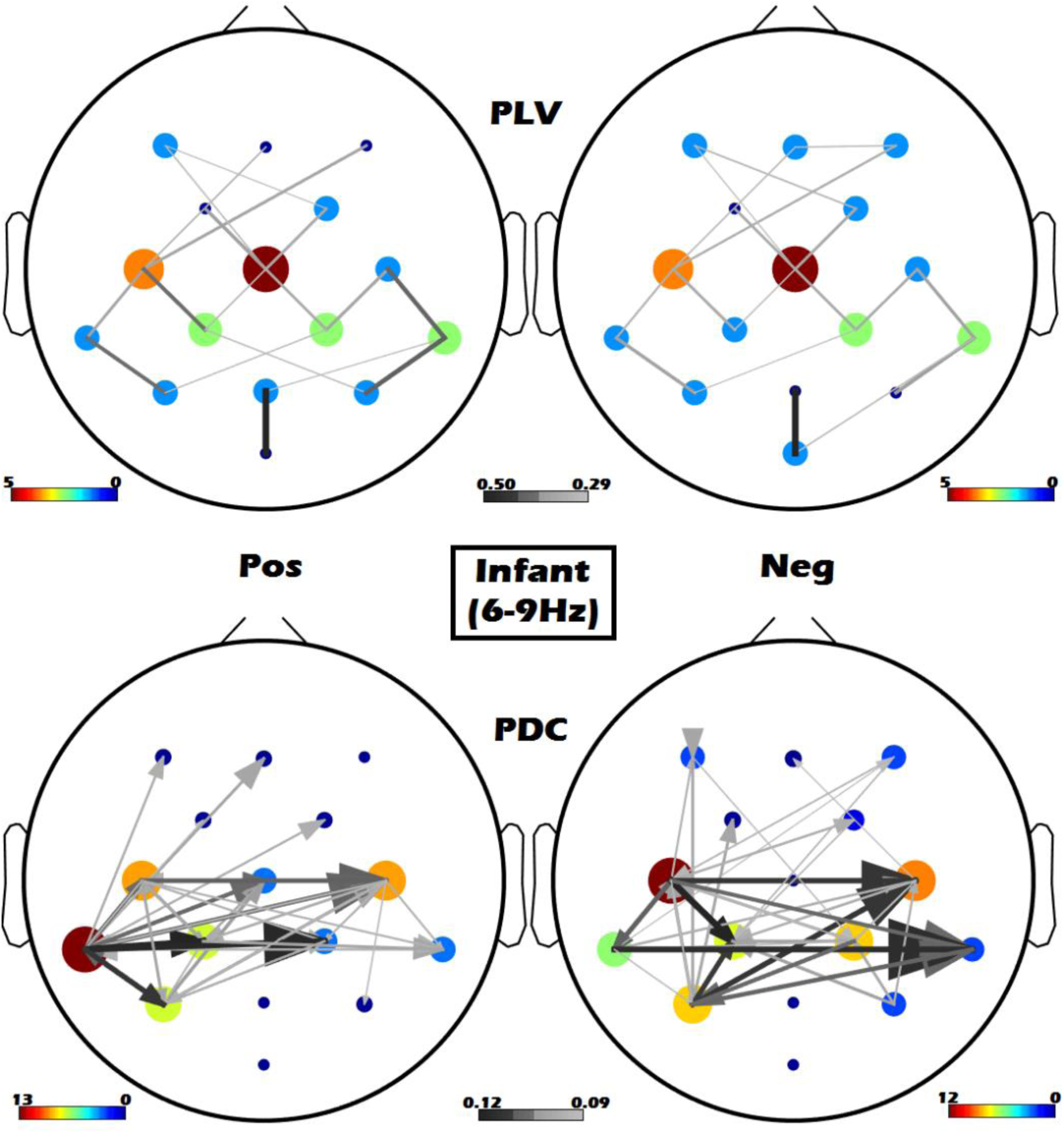
Infant within brain connectivity patterns for the 6-9 Hz band using PLV (top row) and PDC (bottom row). The left column shows the Pos condition and right column shows the Neg condition. For each subplot, the colour and size of each node is proportional its degree, where hotter colours indicate higher values and cooler colours indicate lower values. The weight of the edges in the networks are represented in grey scale, where darker colours indicate stronger connections. For the PDC measure, arrows represent the directionality of connections, ending in the node receiving the information flow.

For adults, significant connections were strongest in temporal-parietal regions for both connectivity metrics (Figure 4). However, whereas PDC-derived networks emphasised interhemispheric (left-right) connections, PLV links frequently connected a node to its closest neighbours, perhaps reflecting volume conduction effects.

Infants’ topographies were characterised by strong connections in central and tempo-parietal regions. Similar to what was observed for adults, infants’ PDC network also showed strong interhemispheric patterns of connectivity. This pattern is consistent with the early emergence of interhemispheric functional connectivity between primary brain regions, which has been demonstrated to exist even in the fetal brain (Anderson & Thomason, 2013; Fransson et al., 2007).

Paired t-tests of intra-brain network density (computed separately for each participant (adult and infant) and metric (PLV and PDC)) revealed that there were no significant differences between *Pos* versus *Neg* conditions, for either metric or participant (mean *Pos-Neg* density: PLV adult = 0.0, *p*=1.00; PLV infant = −0.01, *p*=1.00; PDC adult = 0.02, *p*=0.20; PDC infant = −0.01, p=0.99; *p*-values corrected for multiple comparisons, Tukey HSD).

#### 3.1.2 Intra-brain graph indices

Next we assessed the topology of adults’ and infants’ networks to see if these properties differed across *Pos* and *Neg* experimental conditions. Recall that four graph indices were computed to represent different aspects of network topology for intra-brain measures (individual or basic metrics, measures of segregation, integration and centrality). Table 1 provides a summary of the Strength (S), Global Efficiency (GE), Transitivity (T) and Betweenness Centrality (BC) values obtained for PLV and PDC connectivity measures, for each experimental condition.

**Table 1.** Results of Repeated Measures ANOVAs for (a) Adult and (b) Infant networks assessing the effect of experimental Condition (Pos/Neg) on the four graph indices (strength [S], transitivity [T], global efficiency [GE], and betweenness centrality [BC]) computed from PDC and PLV measures. Statistically significant differences (*p<0.05) are highlighted in bold and shaded. Means differences are calculated as Pos-Neg.

| (a) Adult |  |  |  |  |  |  |
| --- | --- | --- | --- | --- | --- | --- |
| Index | PDC |  |  | PLV |  |  |
|  | F(1,28) | p | Means diff | F(1,28) | p | Means diff |
| Strength (S) | 0.53 | 0.47 | -0.05 | 2.17 | 0.15 | -0.38 |
| Transitivity (T) | 5.87 | **0.02\*** | -0.01 | 0.79 | 0.37 | -0.03 |
| Global Efficiency (GE) | 8.5E-2 | 0.77 | +0.001 | 2.94 | 0.09 | +0.39 |
| Betweenness Centrality (BC) | 0.08 | 0.77 | +0.14 | 6.47 | **0.02\*** | +6.21 |

*Table 1. Results of Repeated Measures ANOVAs for (a) Adult and (b) Infant networks assessing the effect of experimental Condition (Pos/Neg) on the four graph indices (strength [S], transitivity [T], global efficiency [GE], and betweenness centrality [BC]) computed from PDC and PLV measures.
| (b) Infant |  |  |  |  |  |  |
| --- | --- | --- | --- | --- | --- | --- |
| Index | PDC |  |  | PLV |  |  |
|  | F(1,28) | p | Means diff | F(1,28) | p | Means diff |
| Strength (S) | 1.31 | 0.26 | -0.06 | 0.14 | 0.71 | +0.05 |
| Transitivity (T) | 0.94 | 0.34 | -0.004 | 0.007 | 0.93 | +0.002 |
| Global Efficiency (GE) | 1.8E-5 | 0.99 | +0.04 | 0.47 | 0.49 | -0.20 |
| Betweenness Centrality (BC) | 0.01 | 0.90 | -0.37 | 1.40 | 0.25 | -4.03 |

For adults, we observed limited differences in network topology as a function of emotional valence. Namely, PDC Transitivity *decreased* and PLV Betweenness Centrality *increased* (*p*=.02 for both; Hedges’g=-0.861 and 0.936 respectively) for the *Pos* condition with respect to *Neg* condition. For infants however, no statistically significant differences were observed between conditions for any graph metric (*p*>.25 for all indices).

### 3.2 Inter-brain connectivity

#### 3.2.1 Inter-brain connectivity by experimental condition

Figures 6 and 7 show the significant inter-brain connections (relative to surrogate data, see Methods Section 2.8) that were observed between mothers and infants during *Pos* and *Neg* conditions for PLV and PDC metrics respectively.

**Figure 6.**
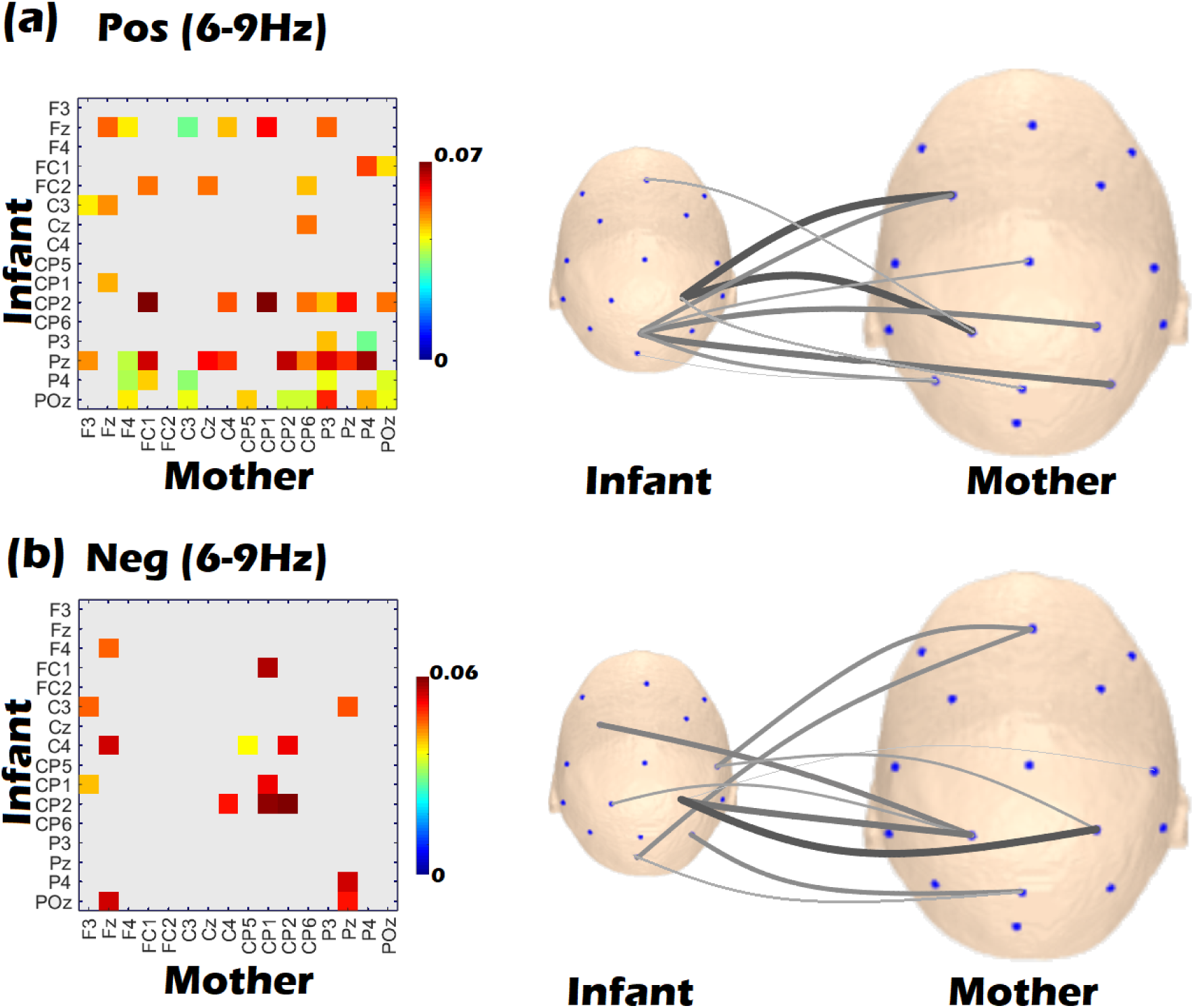
Grand average inter-brain connectivity PLV matrices for Positive (a) and Negative (b) conditions in the 6-9 Hz band. On the left side is the connectivity matrix; rows correspond to infants’ EEG channels and columns correspond to mothers’ channels. Statistically significant inter-brain connections are shown in colour (redder colours indicate higher PLV values) and non-significant connections are shown in light grey. On the right size, topographical head plots of significant inter-brain connections for Pos condition (top) and Neg condition (bottom) are shown. In both cases infants are shown on the left and mothers on the right. The weight of edges in the inter-brain network is represented in grey scale, with darker colours and thicker lines indicating stronger connections. For clarity only the 10 highest connections are plotted in the topographies, whereas in the matrices all significant connections are indicated.

**Figure 7.**
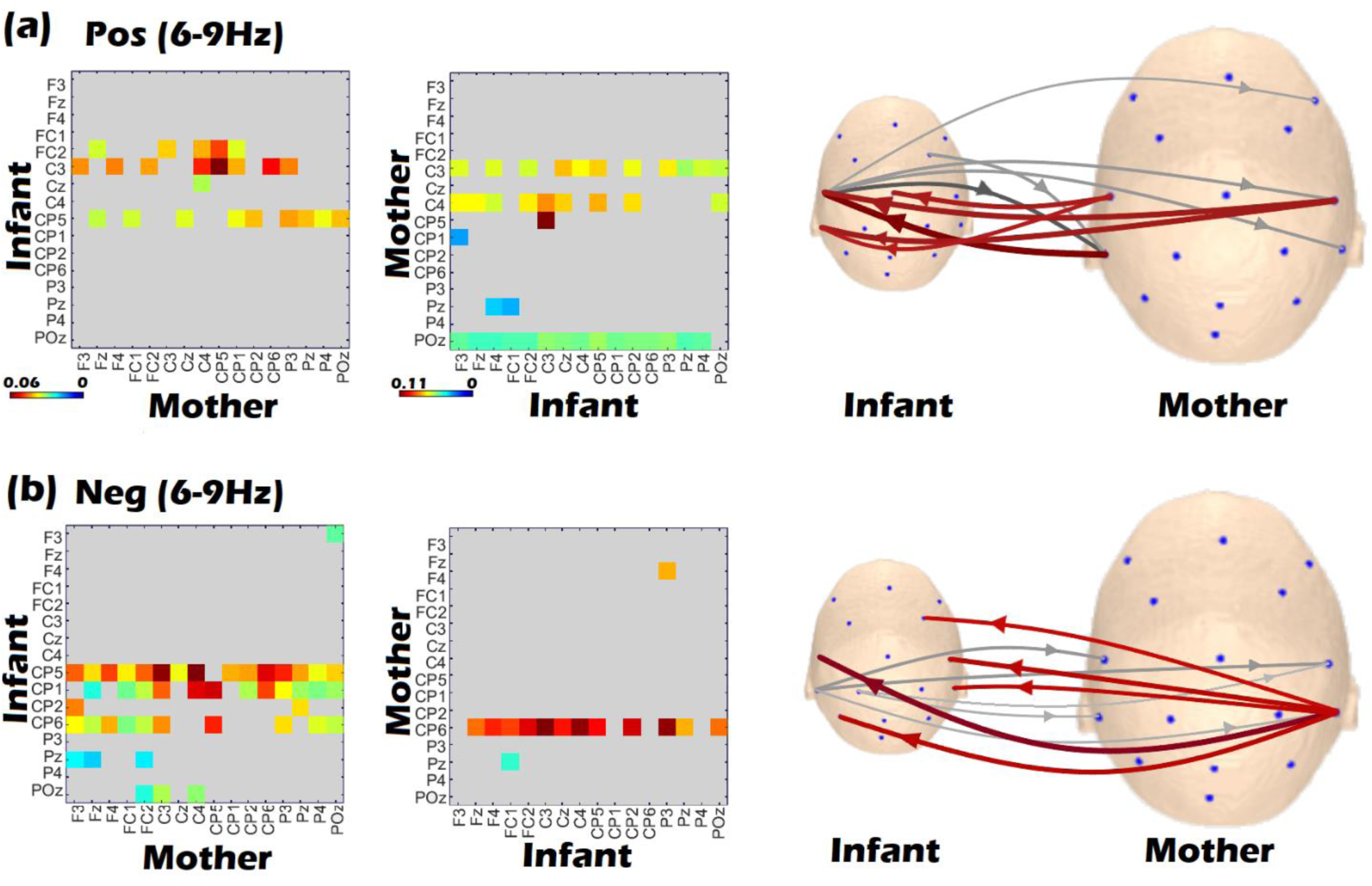
Grand average inter-brain connectivity PDC matrices for Positive (a) and Negative (b) conditions in the 6-9 Hz band. On the left side are the connectivity matrices for each direction of ‘sending’: (infant to mother (left matrix) and mother to infant (right matrix)). Connections which are not statistically significant are marked in grey (redder colours indicate higher PDC values). On the right side (third column) topographical head plots of significant connections for Pos (top) and Neg (bottom) conditions are shown. In both cases infants are shown on the left and mothers on the right. Connections from the infant to the mother are shown in grey whilst connections from the mother to the infant are shown in red. For both directions of sending, darker and thicker lines indicate stronger connections. For clarity only the 5 highest connections in each direction (10 in total) are plotted in the scalp topographies, whereas in the matrices all significant connections are indicated.

For the PLV metric (Figure 6, left), the *Pos* condition (first row) suggested a denser connection between mothers and infants than the *Neg* condition (second row). To statistically assess this difference, we computed the inter-brain density (IBD) of the network in each condition (see Figure 8). The results of the ANOVA indeed revealed a significant effect (F(1,28)=234.09, *p*<.001, mean difference=+0.101), confirming that IBD was significantly higher during the *Pos* than the *Neg* condition.

**Figure 8.**
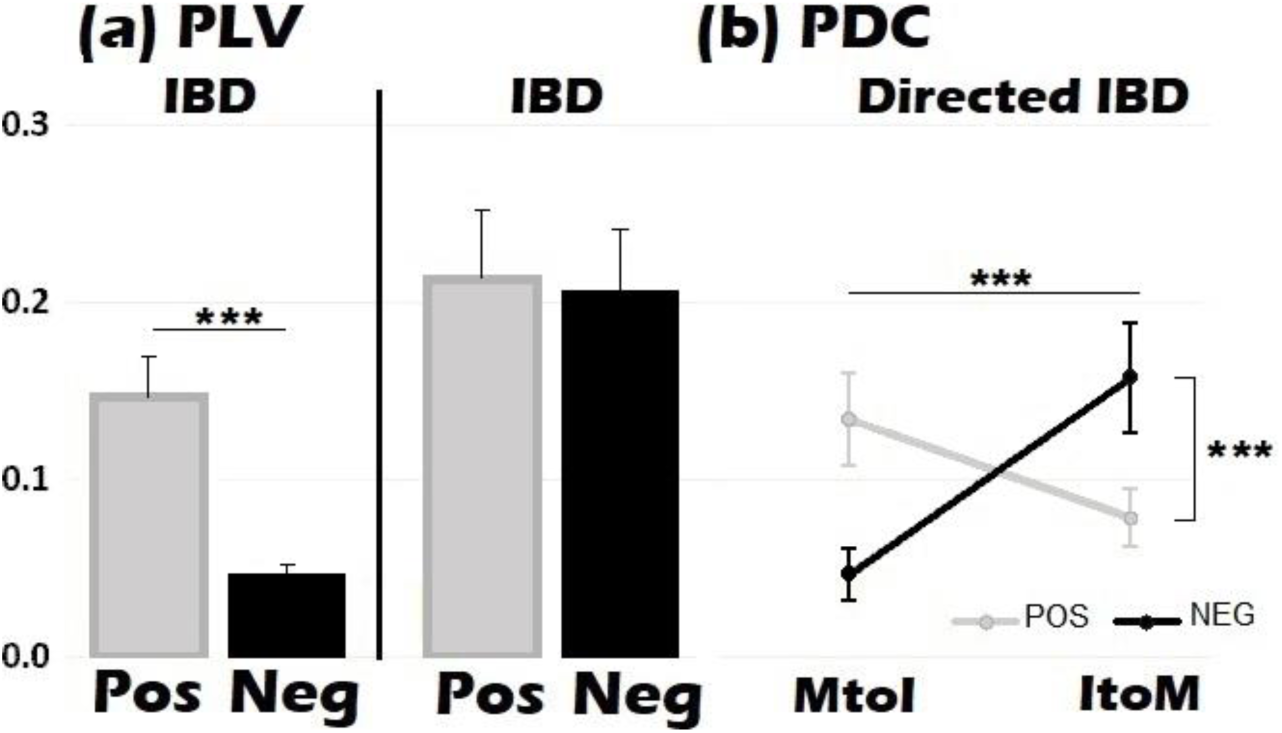
Inter-brain density (IBD) for (a) PLV and (b) PDC. For PDC two different metrics were obtained: total IBD (calculated as for PLV) and directed IBD (mother to infant [MtoI] and infant to mother [ItoM]). ***p<0.001.

A similar IBD analysis was carried out using the PDC metric (Figure 8b), where the inter-brain density for each direction of sending (from infant to mother [ItoM] and from mother to infant [MtoI]) was estimated in addition to the total density (sum of infant to mother and mother to infant IBD). The ANOVA results indicated that for *total* IBD, there was no significant difference between conditions (F(1,28)=0.42, *p*=0.52, mean difference=+0.009). However, analysis of *directed* IBD (Repeated Measures ANOVA with Condition and Direction as within-subjects factors) revealed a significant main effect of sending Direction (F(1,14)=30.05, *p*<.0001, η^2^p = .68) where the ItoM network was more densely connected than the MtoI network overall (Figure 8b, right subplot). Further, a significant interaction was observed between Condition and Direction (F(1,14) = 326.13, *p*<.0001, η^2^p = .96). Post hoc analysis revealed that for MtoI (Mothers ‘sending’ to Infants), inter-brain density was significantly higher for *Pos* > *Neg* (*p*<.001). But for ItoM, inter-brain density was higher for *Neg* > *Pos* (*p*<.001).

#### 3.2.2 Inter-brain graph indices

To quantify topological differences in the pattern of inter-brain connectivity between experimental conditions, two inter-brain graph indices were computed on the thresholded connectivity matrices: Strength and Divisibility (see Section 2.10.2 for full descriptions).

##### Overall Strength

Across both PLV and PDC metrics, Figure 9 shows that the mother-infant inter-brain network had significantly greater strength during the *Pos* condition as compared to the *Neg* condition (PLV *Pos* = 0.096 (±0.026), PLV *Neg* = 0.047 (±0.011); *p*<0.01, Hedges’ g=2.34). The same was true for PDC when both directions of influence (MtoI and ItoM) were averaged (PDC *Pos* = 0.24 (±0.006), PDC *Neg* = 0.17 (±0.07); *p*<.05, Hedges’ g=0.87).

**Figure 9.**
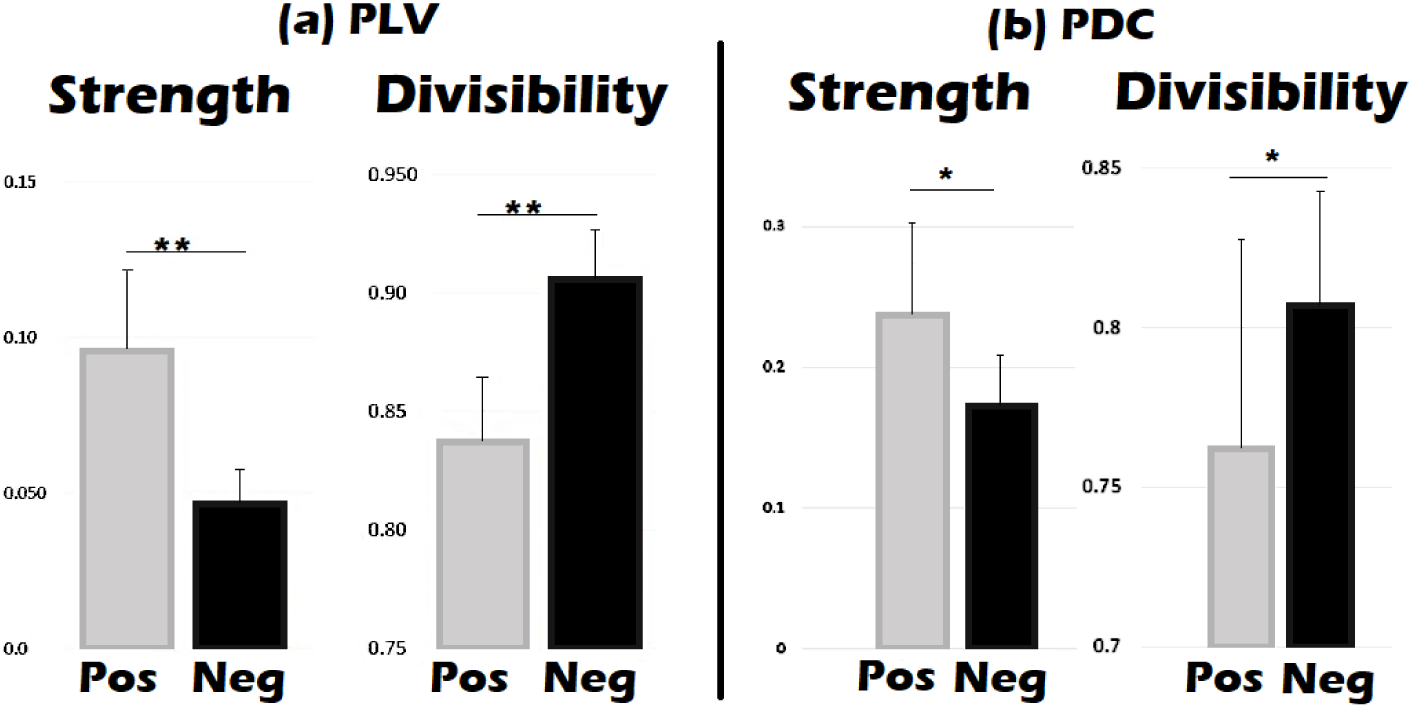
Strength and divisibility inter-brain graph connectivity indices for (a) PLV and (b) PDC, for positive and negative conditions. ** p<0.01, *p<0.05 (false discovery rate corrected)

##### Overall Divisibility

Across both PLV and PDC metrics (see Figure 9), we consistently observed significantly *reduced* divisibility in the *Pos* condition as compared to the *Neg* condition (PLV *Pos*=0.837±0.02, PLV *Neg*=0.906±0.02, *p*<0.01, Hedges’ g = −2.66; PDC *Pos*=0.762±0.04, PDC *Neg*=0.806±0.05, *p*=0.02, Hedges’ g = −0.84). These results indicate greater integration between mothers’ and infants’ sub-networks during the *Pos* condition.

Finally, taking advantage of the property of directionality for the PDC metric, directed Strength and Divisibility were calculated for each participant and condition. Figure 10 shows the average directed strength and directed divisibility for mother to infants (MtoI) and infants to mothers (ItoM), for each condition (*Pos* and *Ne*g). For directed Strength, a Repeated Measures ANOVA revealed a significant main effect of Condition (F(1,14) = 6.56, *p*<.05, η^2^p = .32, Pos> Neg), a significant main effect of Direction (F(1,14) = 11.18, *p*<.01, η^2^p = .44, MtoI > ItoM) and a significant interaction between Condition and Direction (F(1,14)=8.72, *p*<.05, η^2^p = .38). Post hoc analysis of the interaction revealed that whereas MtoI sending was significantly higher for *Pos* > *Neg* (*p*<.01), there was no significant difference between conditions for ItoM sending (*p*=.99).

**Figure 10.**
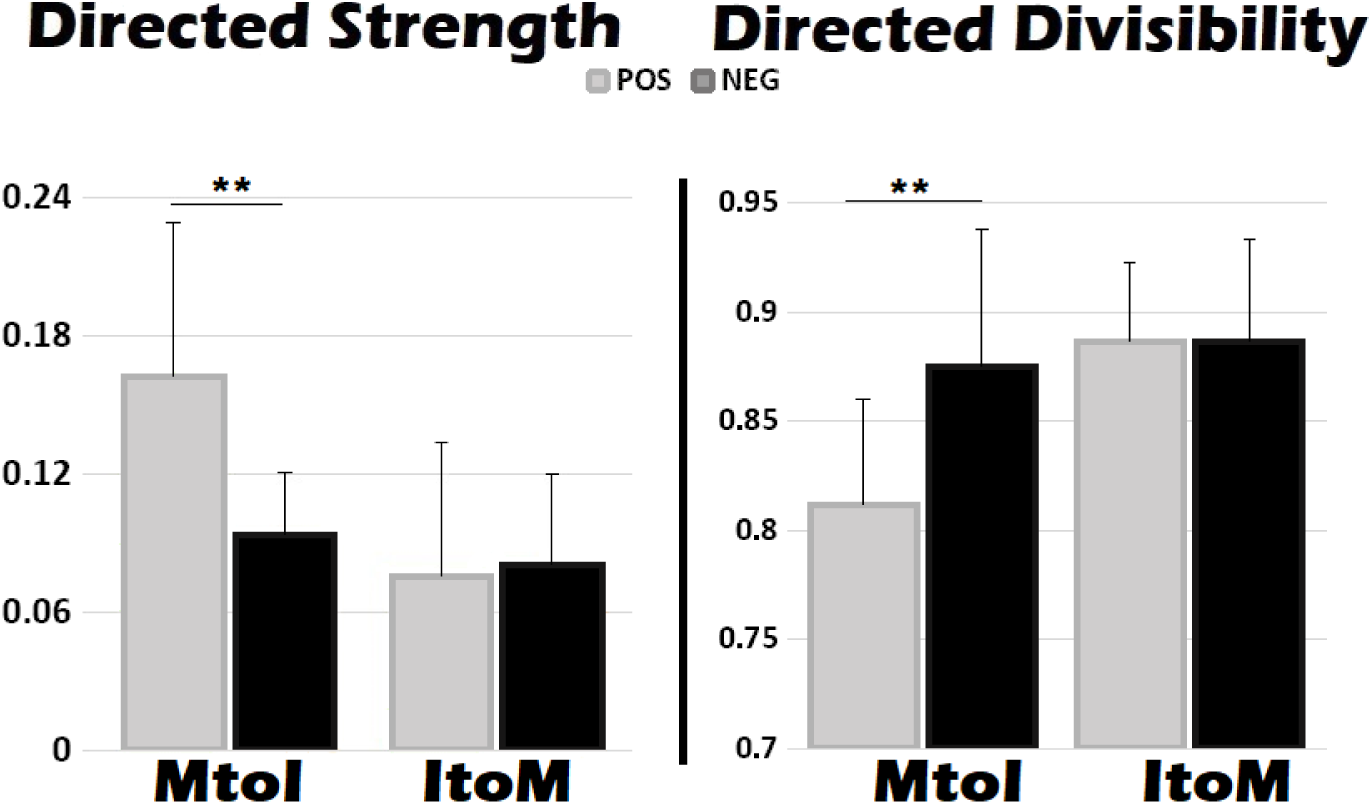
Directed Strength (left) and Divisibility (right) for PDC inter-brain connectivity, for mothers to infants (MtoI) and infants to mothers (ItoM). ** p<0.01

A complementary pattern was observed for directed Divisibility (Figure 10, right). The Repeated Measures ANOVA revealed significant main effects of Condition (F(1,14) = 6.06, *p*<.05, η^2^p = .30, Neg> Pos) and Direction (F(1,14) = 10.50, p<.01, η^2^p = .43, ItoM > MtoI), as well as a significant interaction between Condition and Direction (F(1,14)=7.29, p<.05, η^2^p = .34). This pattern is consistent with the results showed for Strength (where values were higher for the *Pos* condition), as both metrics are inversely related (Ciaramidaro et al., 2018; De Vico et al., 2010; Toppi et al., 2016). Post hoc analysis of the interaction revealed that whereas MtoI divisibility was significantly higher for *Neg* > *Pos* (*p*<.01), there was no significant difference between conditions for ItoM sending (p=.90).

#### 3.2.3 Control for acoustic differences across conditions

Finally, we were concerned that the observed inter-brain connectivity differences between *Pos* and *Neg* conditions could have arisen from sensorimotor differences in the production or perception of *Pos* versus *Neg* maternal utterances, rather than from emotional valence effects per se. Accordingly, we sought to establish (1) whether there were significant differences in the acoustic properties of maternal *Pos* and *Neg* utterances, and if so (2) whether these acoustic differences accounted for our observed results. As reported in the Supplementary Materials (S3), these control analyses showed that the addition of loudness (which differed across conditions) as a covariate in our statistical analyses did not introduce any major systematic changes to the previously-reported results on inter-brain connectivity.

## 4 DISCUSSION

This study aims to describe changes in parent-infant intra-and inter-brain network topology as a function of the valence of emotions displayed by mothers during social interaction with their infants. Social interaction and cooperative communication are of great importance in our daily lives. Previous studies have reported changes in adult-adult interpersonal neural connectivity during cooperative-competitive games (Astolfi et al., 2015; Astolfi et al., 2010; Ciaramidaro et al., 2018; Filho et al., 2016; Sinha et al., 2017), imitation (Delaherche et al., 2015), cooperative action (Müller et al., 2013; Sciaraffa et al., 2017) and verbal spoken communication (Tadić et al., 2016). However, such dyadic neuroimaging studies have usually involved adult participants, and it is not known if, and how, infants’ connectivity with their parents is also modulated by the emotional quality of social interaction.

Here, we find that emotional valence during social interaction (positive or negative) significantly modulates the inter-brain network topology of mother-infant dyads. For both non-directed (phase-locking value, PLV) and directed (partial directed coherence, PDC) measures of connectivity, the inter-brain network showed significantly higher Strength and lower Divisibility for positive as compared to negative emotional states. When considering the *direction* of information flow within the dyad (PDC only), mothers’ influence on and connectedness to their infant was consistently higher during positive than negative emotional states across all directed indices. Conversely, infant-to-parent directed inter-brain density (IBD) was higher during negative emotions, although network Strength and Divisibility showed no significant difference. These results highlight the contrasting role of mothers and infants in modulating the strength and integration of dyadic neural connections during different emotional states. This valence selectivity may be due to infants’ stronger responses to negative than positive maternal affect, which is known to trigger an increase in infants’ own visual scanning and attention solicitation behaviour (Toda & Fogel, 1993; Weinberg et al., 1999; 1996).

By contrast, we observed no emotional valence differences in the *intra*-brain network topology of infants for all graph metrics assessed. Further, intra-brain density did not differ across conditions for either mothers or infants, using both directed and non-directed metrics. However, some emotional valence differences in *maternal* network organisation were observed: maternal Transitivity (for PDC) was decreased and Betweenness Centrality (for PLV) was increased during positive emotions. Betweenness centrality is a measure of the “importance” of each node to the transit of information across the network. Nodes with high Betweenness act as centralised hubs in a network. Hence, if a network has high Betweenness new information can spread more easily throughout the network, facilitating functional integration. This is congruent with lower Transitivity, which is a measure of segregated neural processing. Accordingly, during the communication of positive as compared to negative emotions, maternal neural networks were more strongly integrated, permitting more efficient neural communication (van den Heuvel & Sporns, 2013). It was surprising that no significant differences in infants’ network topology were observed across conditions, for all metrics assessed, especially given that global Alpha network characteristics can be reliably assessed in 10 month infants (Velde et al., 2019). One possible explanation could be that infants’ neural networks for processing positive and negative emotions may (at this point in development) not yet be structurally differentiated, as compared to adults’. For example, previous work has demonstrated that in adults’ brains, structural maturational changes occur during which inefficient connections are pruned to conserve energy (Boersma et al., 2011; Bullmore & Sporns, 2009; Rotem-Kohavi et al., 2017). Further, although structural hubs emerge relatively early during brain development, many of these are still in a relatively immature functional state, with those in visual and motor regions most functionally active (Fransson et al., 2011).

It is important that these null results are not misinterpreted as indicating that there are no differences in neural activation *per se* within the brains of infants with respect to positive and negative emotions. In fact, when we directly contrasted the neural activation levels for individual connections (without considering network organisation or topology), our supplementary analysis revealed extensive activation differences between *Positive* and *Negative* conditions for both mothers and infants (see Supplementary Materials Section S2). These differences in neural activation were observed particularly in terms of hemispheric lateralisation, which is consistent with prior literature (Coan & Allen, 2004; Davidson, 1984, 1998). Rather, our current findings add to the existing literature by showing that emotional valence modulates the topology of the *inter*-brain network (that is, how information flows between mothers’ and infants’ brains) even more strongly than it modulates to the topology of infants’ *intra*-brain network (i.e. how information flows within the infant’s brain).

### 4.1 Limitations

One limitation of the current work is that the study included a relatively small sample size of N=15 dyads. As a result, individual differences in dyadic emotional processing could not be examined. A second limitation is that a semi-naturalistic experimental design was used in order to facilitate social interaction between mothers and their infants. However, the ecological setting increased the complexity of data analysis, for example in terms of the number and variation in myogenic artifacts contained in mothers’ and infants’ EEG data. This necessitated more stringent data rejection and baselining pre-processing steps in order to account for potentially spurious effects arising from these artifacts (see also Section S3 of the Supplementary Materials for an evaluation of the effect of maternal speech acoustics – and by extension, speech articulation effects - on our main results). A final consideration was with regard to how volume conduction effects could have affected our connectivity analyses. For example, we noted that volume conduction effects could have biased infants’ intra-brain network topography as computed by the PLV metric (Section 3.1.1). However, the main comparison of interest here was between experimental conditions. Since volume conduction effects would be expected to affect both conditions in a similar way, we did not expect volume conduction to confound the interpretation of our main results.

### 4.2 Conclusion

Here, we adopted a dual connectivity approach to assess the effect of emotional valence on the topology of the parent-infant joint neural network. We found that inter-brain network indices (density, strength and divisibility) consistently revealed strong effects of emotional valence on the parent-child connection, whereby parent and child showed stronger integration of their neural processes during positive than negative emotional states. By contrast, only weak valence effects were detected for *intra*-brain connectivity. Further, directed inter-brain metrics (PDC) revealed that mothers had a stronger directional influence on the dyadic network during positive emotional states, whereas infants had a stronger influence on the network during negative emotional states. These results suggest that the parent-infant inter-brain network is strongly modulated by the emotional quality and tone of dyadic social interactions, and that inter-brain graph metrics may be successfully applied to elucidate these effects.

## ACKNOWLEDGEMENTS

This research was funded by a UK Economic and Social Research Council (ESRC) Transforming Social Sciences Grant ES/N006461/1 (to V.L. and S.W.), a Nanyang Technological University start-up Grant M4081585.SS0 (to V.L.), and an ESRC Future Research Leaders Fellowship ES/N017560/1 (to S.W.).

The authors declare no competing interests. Anonymised data is available from the authors upon request.

## ETHICS STATEMENT

This study was carried out in accordance with the recommendations of Cambridge Psychology Research Ethics Committee with written informed consent from all subjects. Parents gave written informed consent on behalf of their children in accordance with the Declaration of Helsinki. The protocol was approved by the Cambridge Psychology Research Ethics Committee (PRE.2016.029).

## REFERENCES

Adolphs, R. (2002). Recognizing emotion from facial expressions: psychological and neurological mechanisms. Behavioral and Cognitive Neuroscience Reviews. https://doi.org/10.1177/1534582302001001003

Akaike, H. (1974). A new look at the statistical model identification. IEEE Trans. Automat. Contr., 19, 716–723.

Aktar, E., Mandell, D. J., de Vente, W., Majdandžić, M., Raijmakers, M. E. J., & Bögels, S. M. (2016). Infants’ Temperament and Mothers’, and Fathers’ Depression Predict Infants’ Attention to Objects Paired with Emotional Faces. Journal of Abnormal Child Psychology, 44(5), 975–990. https://doi.org/10.1007/s10802-015-0085-9

Allen, J. J. B., Keune, P. M., Schönenberg, M., & Nusslock, R. (2018). Frontal EEG alpha asymmetry and emotion: From neural underpinnings and methodological considerations to psychopathology and social cognition. Psychophysiology, 55(1), e13028. https://doi.org/10.1111/psyp.13028

Anderson, A. L., & Thomason, M. E. (2013). Functional plasticity before the cradle: A review of neural functional imaging in the human fetus. Neuroscience & Biobehavioral Reviews, 37(9), 2220–2232. https://doi.org/10.1016/j.neubiorev.2013.03.013

Astolfi, L., Cincotti, F., Mattia, D., De Vico Fallani, F., Salinari, S., Vecchiato, G., … Babiloni, F. (2010). Imagining the social brain: multisubjects EEG recordings during the “Chicken’s game.” In Conf. Proc. IEEE Eng. Biol. Soc. (pp. 1734–1737).

Astolfi, L., Cincotti, F., Mattia, D., Marciani, M. G., Baccala, L. A., & Fallani, F. D. V. (2007). Comparison of different cortical connectivity estimators for high-resolution EEG recordings. Hum. Brain Mapp., 28, 143–157.

Astolfi, L., Toppi, J., Casper, C., Freitag, C., Mattia, D., Babiloni, F., … Siniatchkin, M. (2015). Investigating the neural basis of empathy by EEG hyperscanning during a Third Party Punishment. Conf. Proc. IEEE Eng. Biol. Soc., 2015-Novem, 5384–5387.

Astolfi, L., Toppi, J., De Vico Fallani, F., Vecchiato, G., Salinari, S., Mattia, D., … Babiloni, F. (2010). Neuroelectrical hyperscanning measures simultaneous brain activity in humans. Brain Topography, 23(3), 243–256.

Babiloni, C., Vecchio, F., Infarinato, F., Buffo, P., Marzano, N., Spada, D., … Perani, D. (2011). Simultaneous recording of electroencephalographic data in musicians playing in ensemble. Cortex, 47(9), 1082–1090. https://doi.org/10.1016/j.cortex.2011.05.006

Babiloni, F., & Astolfi, L. (2014). Social neuroscience and hyperscanning techniques: Past, present and future. Neuroscience & Biobehavioral Reviews, 44, 76–93. https://doi.org/10.1016/j.neubiorev.2012.07.006

Baccala, L. A., & Sameshima, K. (2001). Partial directed coherence: a new concept in neural structure determination. Biol. Cybern., 84(6), 463–474.

Benjamini, Y., & Yekutieli, D. (2001). The control of the false discovery rate in multiple testing under depencency. The Annals of Statistics, 29(4), 1165–1188.

Betzel, R. F., Satterthwaite, T. D., Gold, J. I., & Bassett, D. S. (2017). Positive affect, surprise, and fatigue are correlates of network flexibility. Scientific Reports, 7(1), 520. https://doi.org/10.1038/s41598-017-00425-z

Boersma, M., Smit, D. J. A., De Bie, H. M. A., Van Baal, G. C. M., Boomsma, D. I., De Geus, E. J. C., … Stam, C. J. (2011). Network analysis of resting state EEG in the developing young brain: Structure comes with maturation. Human Brain Mapping, 32(3), 413–425. https://doi.org/10.1002/hbm.21030

Brooker, B. H., & Donald, M. W. (1980). Contribution of the speech musculature to apparent human EEG asymmetries prior to vocalization. Brain and Language, 9(2), 226–245. https://doi.org/10.1016/0093-934X(80)90143-1

Bullmore, E., & Sporns, O. (2009). Complex brain networks: graph theoretical analysis of structural and functional systems. Nature Reviews Neuroscience, 10(3), 186–198. https://doi.org/10.1038/nrn2575

Ciaramidaro, A., Toppi, J., Casper, C., Freitag, C. M., Siniatchkin, M., & Astolfi, L. (2018). Multiple-Brain Connectivity During Third Party Punishment: an EEG Hyperscanning Study. Scientific Reports, 8(1), 6822. https://doi.org/10.1038/s41598-018-24416-w

Coan, J. A., & Allen, J. J. B. (2004). Frontal EEG asymmetry as a moderator and mediator of emotion. Biological Psychology, 67(1–2), 7–50. https://doi.org/10.1016/j.biopsycho.2004.03.002

Cohn, J. F., & Tronick, E. Z. (1988). Mother-Infant Face-to-Face Interaction: Influence is Bidirectional and Unrelated to Periodic Cycles in Either Partner’s Behavior. Developmental Psychology. https://doi.org/10.1037/0012-1649.24.3.386

Csibra, G., & Gergely, G. (1998). The teleological origins of mentalistic action explanations: A developmental hypothesis. Developmental Science, 1(2), 255–259. https://doi.org/10.1111/1467-7687.00039

Davidson, R. J. (1984). Affect, cognition, and hemispheric specialization. In C. E. Izard, J. Kagan, & R. Zajonc (Eds.), Emotion, Cognition, and Behavior (pp. 320–365). New York: Cambridge University Press.

Davidson, R. J. (1998). Affective Style and Affective Disorders: Perspectives from Affective Neuroscience. Cognition and Emotion. https://doi.org/10.1080/026999398379628

Dawson, G., Klinger, L. G., Panagiotides, H., Hill, D., & Spieker, S. (1992). Frontal Lobe Activity and Affective Behavior of Infants of Mothers with Depressive Symptoms. Child Development, 63(3), 725–737. https://doi.org/10.1111/j.1467-8624.1992.tb01657.x

De Vico Fallani, F., Nicosia, V., Sinatra, R., Astolfi, L., Cincotti, F., Mattia, D., … Babiloni, F. (2010). Defecting or Not Defecting: How to “Read” Human Behavior during Cooperative Games by EEG Measurements. PLoS ONE, 5(12), e14187. https://doi.org/10.1371/journal.pone.0014187

De Vico Fallani, F., Nicosia, V., Sinatra, R., Astolfi, L., Cincotti, F., Mattia, D., … Babiloni, F. (2010). Investigating the neural basis of empathy by EEG hyperscanning during a Third Party Punishment. Brain Topography, 5(3), 5384– 5387.

Delaherche, E., Dumas, G., Nadel, J., & Chetouani, M. (2015). Automatic measure of imitation during social interaction: A behavioral and hyperscanning-EEG benchmark. Pattern Recognition Letters, 66, 118–126. https://doi.org/10.1016/j.patrec.2014.09.002

Deuker, L., Bullmore, E. T., Smith, M., Christensen, S., Nathan, P. J., Rockstroh, B., & Bassett, D. S. (2009). Reproducibility of graph metrics of human brain functional networks. NeuroImage, 47(4), 1460–1468. https://doi.org/10.1016/j.neuroimage.2009.05.035

Diano, M., Tamietto, M., Celeghin, A., Weiskrantz, L., Tatu, M. K., Bagnis, A., … Costa, T. (2017). Dynamic Changes in Amygdala Psychophysiological Connectivity Reveal Distinct Neural Networks for Facial Expressions of Basic Emotions. Scientific Reports. https://doi.org/10.1038/srep45260

Dikker, S., Wan, L., Davidesco, I., Kaggen, L., Oostrik, M., McClintock, J., … Poeppel, D. (2017). Brain-to-Brain Synchrony Tracks Real-World Dynamic Group Interactions in the Classroom. Current Biology, 27(9), 1375–1380. https://doi.org/10.1016/j.cub.2017.04.002

Ding, M., Bressler, S. L., Yang, W., & Liang, H. (2000). Short-window spectral analysis of cortical event-related potentials by adaptive multivariate autoregressive modeling: data preprocessing, model validation, and variability assessment. Biological Cybernetics, 83(1), 35–45. https://doi.org/10.1007/s004229900137

Dumas, G., Lachat, F., Martinerie, J., Nadel, J., & George, N. (2011). From social behaviour to brain synchronization: Review and perspectives in hyperscanning. IRBM, 32(1), 48–53. https://doi.org/10.1016/j.irbm.2011.01.002

Falk, E. B., & Bassett, D. S. (2017). Brain and Social Networks: Fundamental Building Blocks of Human Experience. Trends in Cognitive Sciences, 21(9), 674–690. https://doi.org/10.1016/j.tics.2017.06.009

Feinman, S. (1982). Social Referencing in Infancy. Merrill-Palmer Quarterly, 28(4), 445–470. https://doi.org/10.2307/23086154

Feinman, S., & Lewis, M. (1983). Social Referencing at Ten Months: A Second-Order Effect on Infants’Responses to Strangers. Child Development, 54(4), 878–887. https://doi.org/10.1111/j.1467-8624.1983.tb00509.x

Feinman, S., & Roberts, D. (1986). The effect of social referencing on 12-month-olds’ responses to a stranger’s attempts to “make friends.” Infant Behavior and Development, 9, 119. https://doi.org/10.1016/S0163-6383(86)80121-7

Feinman, S., Roberts, D., Hsieh, K.-F., Sawyer, D., & Swanson, D. (1992). A Critical Review of Social Referencing in Infancy. In Social Referencing and the Social Construction of Reality in Infancy (pp. 15–54). Boston, MA: Springer US. https://doi.org/10.1007/978-1-4899-2462-9_2

Feldman, R. (2007). Parent-infant synchrony and the construction of shared timing; physiological precursors, developmental outcomes, and risk conditions. Journal of Child Psychology and Psychiatry and Allied Disciplines, 48(3–4), 329–354.

Field, T., Fox, N. A., Pickens, J., & Nawrocki, T. (1995). Relative Right Frontal EEG Activation in 3-to 6-Month-Old Infants of “Depressed” Mothers. Developmental Psychology, 31, 358–363. https://doi.org/10.1037/0012-1649.31.3.358

Field, T., Pickens, J., Fox, N. A., Nawrocki, T., & Gonzalez, J. (1995). Vagal tone in infants of depressed mothers. Development and Psychopathology, 7, 227–231. https://doi.org/10.1017/S0954579400006465

Figuera Jaimes, R., Arellano Ferro, A., Bramich, D. M., Giridhar, S., & Kuppuswamy, K. (2013). Variable stars in the globular cluster NGC 7492. Astronomy & Astrophysics, 556, A20. https://doi.org/10.1051/0004-6361/201220824

Filho, E., Bertollo, M., Tamburro, G., Schinaia, L., Chatel-Goldman, J., di Fronso, S., … Comani, S. (2016). Hyperbrain features of team mental models within a juggling paradigm: a proof of concept. PeerJ, 4, e2457. https://doi.org/10.7717/peerj.2457

Fransson, P., Åden, U., Blennow, M., & Lagercrantz, H. (2011). The functional architecture of the infant brain as revealed by resting-state fMRI. Cerebral Cortex. https://doi.org/10.1093/cercor/bhq071

Fransson, P., Skiold, B., Horsch, S., Nordell, A., Blennow, M., Lagercrantz, H., & Aden, U. (2007). Resting-state networks in the infant brain. Proceedings of the National Academy of Sciences, 104(39), 15531–15536. https://doi.org/10.1073/pnas.0704380104

Garrison, K. A., Scheinost, D., Finn, E. S., Shen, X., & Constable, R. T. (2015). The (in)stability of functional brain network measures across thresholds. NeuroImage, 118, 651–661. https://doi.org/10.1016/j.neuroimage.2015.05.046

Georgieva, S., Lester, S., Yilmaz, M., Wass, S., & Leong, V. (2017). Topographical and spectral signatures of infant and adult movement artifacts in naturalistic EEG. BioRxiv. https://doi.org/10.1109/LED.2010.2052778

Goncharova, I.., McFarland, D. J., Vaughan, T.., & Wolpaw, J. R. (2003). EMG contamination of EEG: spectral and topographical characteristics. Clinical Neurophysiology, 114(9), 1580–1593. https://doi.org/10.1016/S1388-2457(03)00093-2

Gotlib, I. H., Ranganath, C., & Rosenfeld, J. P. (1998). Frontal EEG Alpha Asymmetry, Depression, and Cognitive Functioning. Cognition and Emotion. https://doi.org/10.1080/026999398379673

Goulden, N., McKie, S., Thomas, E. J., Downey, D., Juhasz, G., Williams, S. R., … Elliott, R. (2012). Reversed Frontotemporal Connectivity During Emotional Face Processing in Remitted Depression. Biological Psychiatry, 72(7), 604–611. https://doi.org/10.1016/j.biopsych.2012.04.031

Granger, C. W. J. (1969). Investigating Causal Relations by Econometric Models and Cross-spectral Methods. Econometrica, 37(3), 424–438. https://doi.org/10.2307/1912791

Gunnar, M. R., & Stone, C. (1984). The Effects of Positive Maternal Affect on Infant Responses to Pleasant, Ambiguous, and Fear-Provoking Toys. Child Development, 55(4), 1231. https://doi.org/10.2307/1129992

Gupta, R., ur Rehman Laghari, K., & Falk, T. H. (2016). Relevance vector classifier decision fusion and EEG graph-theoretic features for automatic affective state characterization. Neurocomputing, 174, 875–884. https://doi.org/10.1016/j.neucom.2015.09.085

Hasson, U., Ghazanfar, A. A., Galantucci, B., Garrod, S., & Keysers, C. (2012). Brain-to-brain coupling: a mechanism for creating and sharing a social world. Trends in Cognitive Sciences, 16(2), 114–121. https://doi.org/10.1016/j.tics.2011.12.007

He, B., Dai, Y., Astolfi, L., Babiloni, F., Yuan, H., & Yang, L. (2011). eConnectome: A MATLAB toolbox for mapping and imaging of brain functional connectivity. Journal of Neuroscience Methods, 195(2), 261–269. https://doi.org/10.1016/j.jneumeth.2010.11.015

Hirshberg, L. M., & Svejda, M. (1990). When Infants Look to Their Parents: I. Infants’ Social Referencing of Mothers Compared to Fathers. Child Development, 61(4), 1175–1186. https://doi.org/10.1111/j.1467-8624.1990.tb02851.x

Hoehl, S. (2013). Emotion Processing in Infancy. In H. L. K. Lagattuta (Ed.), Children and Emotion New Insights into Developmental Affective Science. (pp. 1–12). Karger Publishers. https://doi.org/10.1159/000354346

Hoehl, S., Reid, V., & Parise, E. (2019). The Biological Basis of Social Cognition During Development. Neuropsychologia, 126(March), 1–2.

Hoehl, S., & Striano, T. (2008). Neural processing of eye gaze and threat-related emotional facial expressions in infancy. Child Development, 79(6), 1752–1760. https://doi.org/10.1111/j.1467-8624.2008.01223.x

Hoehl, S., & Striano, T. (2010). Infants’ neural processing of positive emotion and eye gaze. Social Neuroscience, 5(1), 30–39. https://doi.org/10.1080/17470910903073232

Hoehl, S., Wiese, L., & Striano, T. (2008). Young Infants’ Neural Processing of Objects Is Affected by Eye Gaze Direction and Emotional Expression. PLoS ONE, 3(6), e2389. https://doi.org/10.1371/journal.pone.0002389

Hornik, R., Risenhoover, N., & Gunnar, M. R. (1987). The Effects of Maternal Positive, Neutral, and Negative Affective Communications on Infant Responses to New Toys. Child Development, 58(4), 937. https://doi.org/10.2307/1130534

Jaffe, J., Beebe, B., Feldstein, S., Crown, C. L., & Jasnow, M. D. (2001). Rhythms of dialogue in infancy: coordinated timing in development. Monographs of the Society for Research in Child Development, 66(2), i-viii,1-149.

Jiang, J., Chen, C., Dai, B., Shi, G., Ding, G., Liu, L., & Lu, C. (2015). Leader emergence through interpersonal neural synchronization. Proceedings of the National Academy of Sciences, 112(14), 4274–4279. https://doi.org/10.1073/pnas.1422930112

Jiang, J., Dai, B., Peng, D., Zhu, C., Liu, L., & Lu, C.-F. (2012). Neural Synchronization during Face-to-Face Communication. Journal of Neuroscience, 32(45), 16064–16069. https://doi.org/10.1523/JNEUROSCI.2926-12.2012

Kabbara, A., Eid, H., El-Falou, W., Khalil, M., Wendling, F., & Hassan, M. (2018). Reduced integration and improved segregation of functional brain networks in Alzheimer’s disease. J. Neural Eng., 15(2), 026023.

Kaye, K., & Fogel, A. (1980). The temporal structure of face-to-face communication between mothers and infants. Developmental Psychology, 16(5), 454–464. https://doi.org/10.1037/0012-1649.16.5.454

Kotz, S. A., Meyer, M., Alter, K., Besson, M., Von Cramon, D. Y., & Friederici, A. D. (2003). On the lateralization of emotional prosody: An event-related functional MR investigation. Brain and Language. https://doi.org/10.1016/S0093-934X(02)00532-1

Lachaux, J.-P., Rodriguez, E., Martinerie, J., & Varela, F. J. (1999). Measuring phase synchrony in brain signals. Human Brain Mapping, 8(4), 194–208. https://doi.org/10.1002/(SICI)1097-0193(1999)8:4<194::AID-HBM4>3.0.CO;2-C

Leong, V., Byrne, E., Clackson, K., Georgieva, S., Lam, S., & Wass, S. V. (2017). Speaker gaze increases information coupling between infant and adult brains. Proceedings of the National Academy of Sciences, 114(50), 13290–13295. https://doi.org/10.1073/pnas.1702493114

Leppänen, J. M., Moulson, M. C., Vogel-Farley, V. K., & Nelson, C. A. (2007). An ERP study of emotional face processing in the adult and infant brain. Child Development, 78(1), 232–245. https://doi.org/10.1111/j.1467-8624.2007.00994.x

Leppänen, J. M., & Nelson, C. A. (2006). The development and neural bases of facial emotion recognition. Advances in Child Development and Behavior, 34, 207–246.

Leppänen, J. M., & Nelson, C. A. (2009). Tuning the developing brain to social signals of emotions. Nature Reviews Neuroscience, 10(1), 37–47. https://doi.org/10.1038/nrn2554

Leppänen, J., Peltola, M. J., Mäntymaa, M., Koivuluoma, M., Salminen, A., & Puura, K. (2010). Cardiac and behavioral evidence for emotional influences on attention in 7-month-old infants. International Journal of Behavioral Development, 34(6), 547–553. https://doi.org/10.1177/0165025410365804

Lu, Q., Li, H., Luo, G., Wang, Y., Tang, H., Han, L., & Yao, Z. (2012). Impaired prefrontal–amygdala effective connectivity is responsible for the dysfunction of emotion process in major depressive disorder: A dynamic causal modeling study on MEG. Neuroscience Letters, 523(2), 125–130. https://doi.org/10.1016/j.neulet.2012.06.058

Marshall, P. J., Bar-Haim, Y., & Fox, N. A. (2002). Development of the EEG from 5 months to 4 years of age. Clinical Neurophysiology, 113(8), 1199–1208. https://doi.org/10.1016/S1388-2457(02)00163-3

Müller, V., Sänger, J., & Lindenberger, U. (2013). Intra- and Inter-Brain Synchronization during Musical Improvisation on the Guitar. PLoS ONE, 8(9), e73852. https://doi.org/10.1371/journal.pone.0073852

Nelson, C. A., & De Haan, M. (1996). Neural Correlates of Infants’ Visual Responsiveness to Facial Expressions of Emotion. Developmental Psychobiology. https://doi.org/10.1002/(SICI)1098-2302(199611)29:7<577::AID-DEV3>3.0.CO;2-R

Nelson, C. A., Morse, P. A., & Leavitt, L. A. (1979). Recognition of facial expressions by seven-month-old infants. Child Development. https://doi.org/10.1111/j.1467-8624.1979.tb02493.x

Newman, M. E. J. (2003). The Structure and Function of Complex Networks. SIAM Review, 45(2), 167–256. https://doi.org/10.1137/S003614450342480

Nichols, T. E., Das, S., Eickhoff, S. B., Evans, A. C., Glatard, T., Hanke, M., … Yeo, B. T. T. (2017). Best practices in data analysis and sharing in neuroimaging using MRI. Nature Neuroscience, 20(3), 299–303. https://doi.org/10.1038/nn.4500

Nicholson, A. A., Friston, K. J., Zeidman, P., Harricharan, S., McKinnon, M. C., Densmore, M., … Lanius, R. A. (2017). Dynamic causal modeling in PTSD and its dissociative subtype: Bottom–up versus top–down processing within fear and emotion regulation circuitry. Human Brain Mapping. https://doi.org/10.1002/hbm.23748

Niso, G., Bruña, R., Pereda, E., Gutiérrez, R., Bajo, R., Maestú, F., & Del-Pozo, F. (2013). HERMES: Towards an integrated toolbox to characterize functional and effective brain connectivity. Neuroinformatics, 11(4), 405–434. https://doi.org/10.1007/s12021-013-9186-1

Ochsner, K. N., Ray, R. R., Hughes, B., Mcrae, K., Cooper, J. C., Weber, J., … Gross, J. J. (2009). Bottom-up and top-down processes in emotion generation: Common and distinct neural mechanisms. Psychological Science. https://doi.org/10.1111/j.1467-9280.2009.02459.x

Péron, J., Grandjean, D., Le Jeune, F., Sauleau, P., Haegelen, C., Drapier, D., … Vérin, M. (2010). Recognition of emotional prosody is altered after subthalamic nucleus deep brain stimulation in Parkinson’s disease. Neuropsychologia. https://doi.org/10.1016/j.neuropsychologia.2009.12.003

Philips, G. R., Daly, J. J., & Principe, J. C. (2017). Topographical measures of functional connectivity as biomarkers for post-stroke motor recovery. Journal of Neural Engineering and Rehabilitation, 14(1).

Reindl, V., Gerloff, C., Scharke, W., & Konrad, K. (2018). Brain-to-brain synchrony in parent-child dyads and the relationship with emotion regulation revealed by fNIRS-based hyperscanning. NeuroImage, 178, 493–502. https://doi.org/10.1016/j.neuroimage.2018.05.060

Rogoff, B. (1990). Apprenticeship in thinking: Cognitive dvelopment in social contexts. *New York*. Oxford University Press.

Rotem-Kohavi, N., Oberlander, T. F., & Virji-Babul, N. (2017). Infants and adults have similar regional functional brain organization for the perception of emotions. Neuroscience Letters, 650, 118–125. https://doi.org/10.1016/j.neulet.2017.04.031

Rubinov, M., & Sporns, O. (2010). Complex network measures of brain connectivity: Uses and interpretations. NeuroImage, 52(3), 1059–1069. https://doi.org/10.1016/j.neuroimage.2009.10.003

Sänger, J., Müller, V., & Lindenberger, U. (2012). Intra- and interbrain synchronization and network properties when playing guitar in duets. Frontiers in Human Neuroscience, 6, 312. https://doi.org/10.3389/fnhum.2012.00312

Sänger, J., Müller, V., & Lindenberger, U. (2013). Directionality in hyperbrain networks discriminates between leaders and followers in guitar duets. Frontiers in Human Neuroscience, 7. https://doi.org/10.3389/fnhum.2013.00234

Sato, W., Kochiyama, T., Uono, S., Yoshikawa, S., & Toichi, M. (2017). Direction of Amygdala-Neocortex Interaction During Dynamic Facial Expression Processing. Cerebral Cortex (New York, N.Y.: 1991), 27(3), 1878–1890. https://doi.org/10.1093/cercor/bhw036

Schilbach, L., Timmermans, B., Reddy, V., Costall, A., Bente, G., Schlicht, T., & Vogeley, K. (2013). Toward a second-person neuroscience. Behavioral and Brain Sciences, 36(4), 393–414. https://doi.org/10.1017/s0140525x12000660

Schwarz, G. (1978). Estimating the dimension of a model. The Annals of Statistics, 6(2), 461–464.

Sciaraffa, N., Borghini, G., Aricò, P., Di Flumeri, G., Colosimo, A., Bezerianos, A., … Babiloni, F. (2017). Brain interaction during cooperation: Evaluating local properties of multiple-brain network. Brain Sciences, 7(7). https://doi.org/10.3390/brainsci7070090

Seth, A. K. (2010). A MATLAB toolbox for Granger causal connectivity analysis. Journal of Neuroscience Methods, 186(2), 262–273. https://doi.org/10.1016/j.jneumeth.2009.11.020

Sinha, N., Maszczyk, T., Zhang, W., Tan, J., & Dauwels, J. (2017). EEG hyperscanning study of inter-brain synchrony during cooperative and competitive interaction. 2016 IEEE International Conference on Systems, Man, and Cybernetics, SMC 2016 - Conference Proceedings, 4813–4818. https://doi.org/10.1109/SMC.2016.7844990

Skoranski, A. M., Lunkenheimer, E., & Lucas-Thompson, R. G. (2017). The effects of maternal respiratory sinus arrhythmia and behavioral engagement on mother-child physiological coregulation. Developmental Psychobiology. https://doi.org/10.1002/dev.21543

Sorce, J. F., Emde, R. N., Campos, J. J., & Klinnert, M. D. (1985). Maternal emotional signaling: Its effect on the visual cliff behavior of 1-year-olds. Developmental Psychology, 21(1), 195–200. https://doi.org/10.1037/0012-1649.21.1.195

Spangler, G. (1991). The emergence of adrenocortical circadian function in newborns and infants and its relationship to sleep, feeding and maternal adrenocortical activity. Early Human Development, 25(3), 197–208. https://doi.org/10.1016/0378-3782(91)90116-K

Tadić, B., Andjelković, M., Boshkoska, B. M., & Levnajić, Z. (2016). Algebraic topology of multi-brain connectivity networks reveals dissimilarity in functional patterns during spoken communications. PLoS ONE, 11(11), 1–25. https://doi.org/10.1371/journal.pone.0166787

Toda, S., & Fogel, A. (1993). Infant response to the still-face situation at 3 and 6 months. Developmental Psychology, 29(3), 532–538. https://doi.org/10.1037/0012-1649.29.3.532

Toppi, J., Borghini, G., Petti, M., He, E. J., De Giusti, V., He, B., … Babiloni, F. (2016). Investigating cooperative behavior in ecological settings: An EEG hyperscanning study. PLoS ONE, 11(4), 1–26.

Toppi, J., De Vico Fallani, F., Vecchiato, G., Maglione, A. G., Cincotti, F., Mattia, D., … Astolfi, L. (2012). How the Statistical Validation of Functional Connectivity Patterns Can Prevent Erroneous Definition of Small-World Properties of a Brain Connectivity Network. Computational and Mathematical Methods in Medicine, 2012, 1–13. https://doi.org/10.1155/2012/130985

van den Heuvel, M. P., & Sporns, O. (2013). An Anatomical Substrate for Integration among Functional Networks in Human Cortex. Journal of Neuroscience. https://doi.org/10.1523/jneurosci.2128-13.2013

Velde B. van der, Haartsen, R., & Kemmer, C. (2019). Test-retest reliability of EEG network characteristics in infants. Brain and Behavior, e01269.

Walden, T. A., & Ogan, T. A. (1988). The Development of Social Referencing. Child Development, 59(5), 1230–1240. https://doi.org/10.1111/j.1467-8624.1988.tb01492.x

Weinberg, M. K., & Tronick, E. Z. (1996). Infant Affective Reactions to the Resumption of Maternal Interaction after the Still-Face. Child Development, 67(3), 905–914. https://doi.org/10.1111/j.1467-8624.1996.tb01772.x

Weinberg, M. K., Tronick, E. Z., Cohn, J. F., & Olson, K. L. (1999). Gender differences in emotional expressivity and self-regulation during early infancy. Developmental Psychology, 35(1), 175–188. https://doi.org/10.1037/0012- 1649.35.1.175

Whitham, E. M., Pope, K. J., Fitzgibbon, S. P., Lewis, T., Clark, C. R., Loveless, S., … Willoughby, J. O. (2007). Scalp electrical recording during paralysis: Quantitative evidence that EEG frequencies above 20 Hz are contaminated by EMG. Clinical Neurophysiology, 118(8), 1877–1888. https://doi.org/10.1016/j.clinph.2007.04.027

