## Supplementary Materials for "Emotional valence modulates the topology of the parent-infant inter-brain network"

### S.1. Thresholding effects

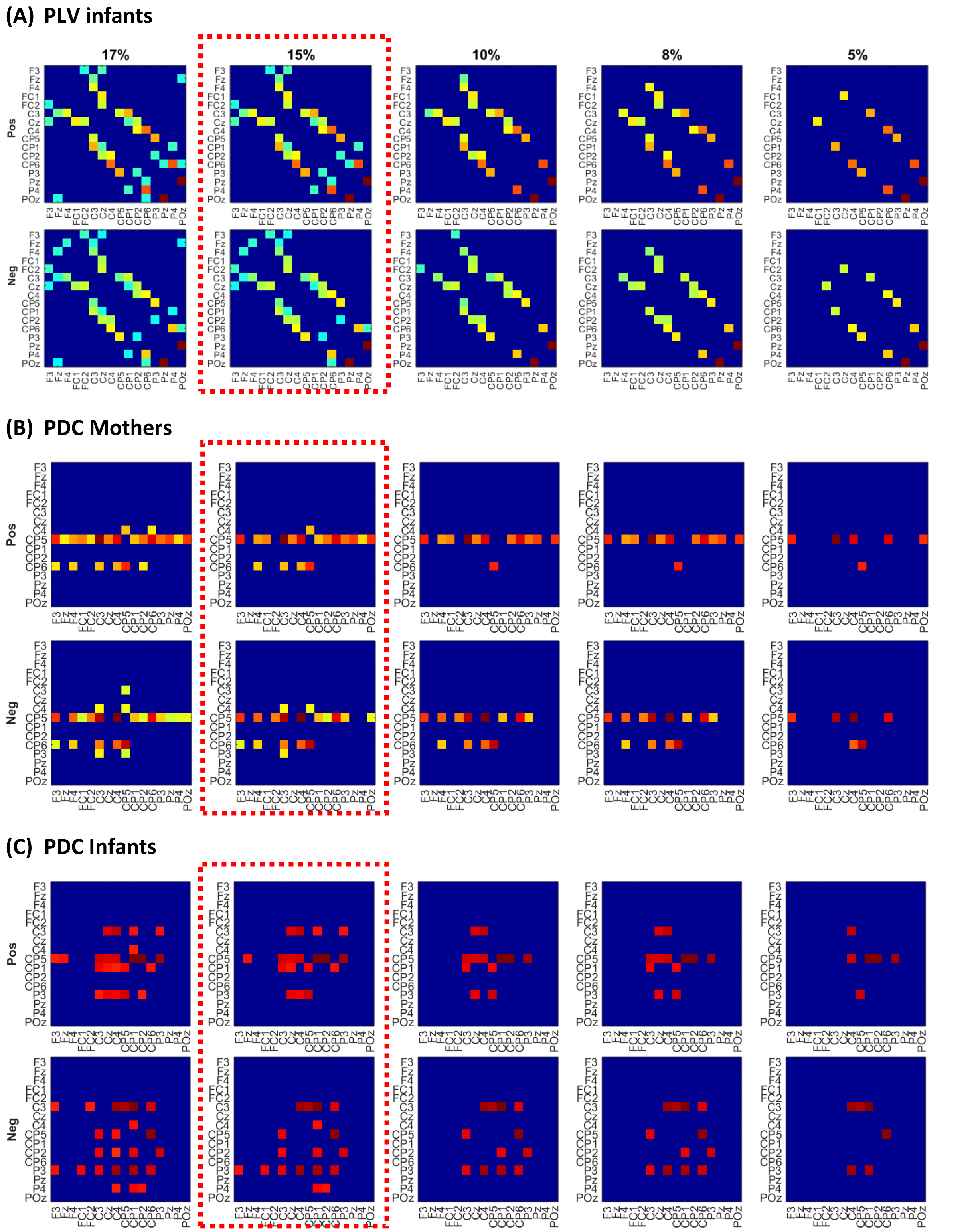


*Figure S1. Effect of applying different thresholds to the infant and adult data for each condition (Pos and Neg) and connectivity metric (PLV and PDC). A series of thresholds ranging from 17% (left column) to 5% (right) column is applied. The dashed rectangle indicates the selected threshold (15%) used in the current analysis.*

### S.2. Intra-brain activation across conditions

To directly contrast neural activation between *Pos* and *Neg* conditions, two-sample t-tests were performed using the statistically validated (but non-thresholded) individual connections for PLV and PDC measures. Trials where at least one condition value (*Pos* or *Neg*) was not significantly above chance were excluded (yielding different numbers of number of trials across conditions). A significance level of 5% was used, posteriorly adjusted for multiple comparisons using the false discovery rate (FDR) procedure (Benjamini & Yekutieli, 2001). Figures S2 and S3 show the results for adults and infants respectively in the 6-9 Hz band using PLV (top row) and PDC (bottom row).

Mothers showed predominantly stronger activation for the *Neg* condition than the *Pos* condition across both PLV and PDC connectivity metrics. In the scalp plots of Figure S2, this may be observed as a larger number of dark green (*Neg*>*Pos*) than light green (*Pos*>*Neg*) connections. These differences were widely distributed across all scalp regions and could reflect the increased arousal (i.e. stress) and cognitive effort that was required by mothers to model a negative emotion to their infants.

By contrast, infants showed more restricted differences between *Pos* and *Neg* conditions (see Figure S3). First, for both PLV and PDC metrics, *increased* activation for the *Pos* condition was observed over the *left* parieto-occipital region, which is consistent with prior reports on left-hemisphere processing of positive emotions (Coan & Allen, 2004; Davidson, 1984). By contrast, stronger activation for the *Neg* condition was observed in the posterior occipital and central scalp regions. We also observed a larger number of significantly-different connections for the PDC metric (32 significant connections) as compared to the PLV metric (13 significant connections) for the infant data. This difference could reflect the sensitivity of the PDC measure to changes in both EEG power and phase, whereas the PLV measure detects only phase differences.

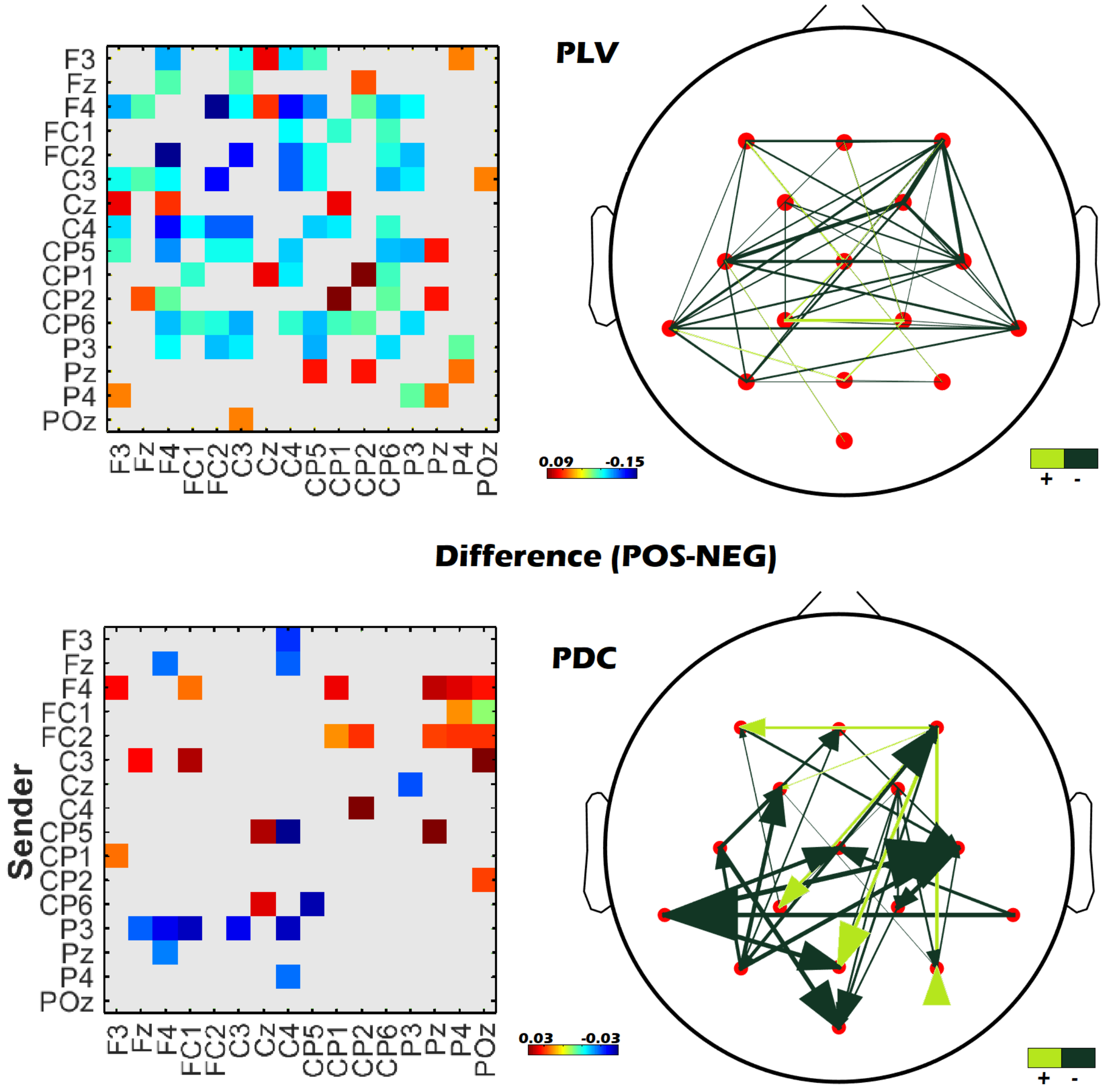


*Figure S2. Adult neural activation differences between Pos and Neg conditions using PLV (top row) and PDC (bottom row) metrics for the 6-9 Alpha band. Both representations (matrices and topographies) show only statistically significant (p<0.5) differences between conditions (Pos minus Neg). In the matrices hotter colours represent stronger connections, cooler colours represent lower values. The head plots show in light green links where Pos values are higher than Neg values, hence, the difference is positive. Contrarily, dark green links are stronger for the Neg condition than the Pos condition.*

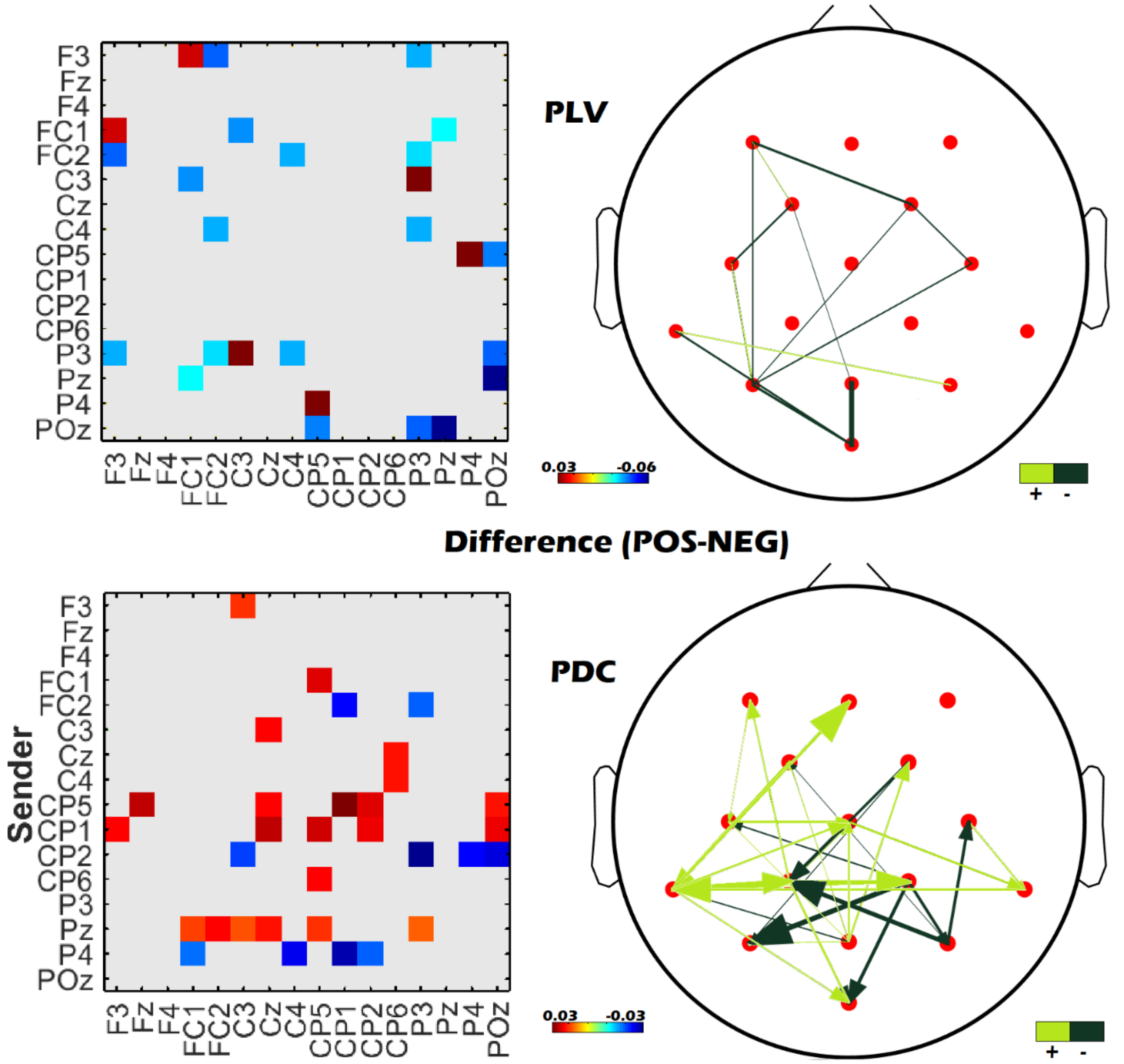


*Figure S3. Infant neural activation differences between Pos and Neg conditions using PLV (top row) and PDC (bottom row) metrics for the 6-9 Alpha band. Both representations (matrices and topographies) show only statistically significant ($p<0.5$) differences between conditions (Pos minus Neg). In the matrices hotter colours represent stronger connections, cooler colours represent lower values. The head plots show in light green links where Pos values are higher than Neg values, hence, the difference is positive. Contrarily, dark green links are stronger for the Neg condition than the Pos condition.*

### S.3. Effect of maternal speech acoustic parameters on connectivity analyses

Here, we assess whether the parent-infant inter-brain network is significantly modulated by the emotional valence of social interaction. However, it is possible that any observed neural connectivity differences across *Pos* and *Neg* conditions may in fact arise from sensorimotor differences in the production or perception of *Pos* versus *Neg* maternal utterances. Accordingly, we sought to establish (1) whether there were significant differences in the acoustic properties of maternal *Pos* and *Neg* utterances, and if so (2) whether these acoustic differences accounted for our observed results. We addressed the second aim by conducting ANCOVA analyses, where the relevant acoustic parameters were added as covariates to the original analyses. Due to technical difficulties, measurements of pitch and loudness could not be obtained from two out of the fifteen dyads, thus the following analyses are based on data from thirteen (87%) dyads. As reported in detail below, the addition of loudness (which differed across conditions) as a covariate in our statistical analyses did not produce any major systematic changes to the main results on inter-brain connectivity.

#### a) Acoustic differences across conditions

As reported in the main text, maternal speech duration did not differ significantly across conditions ($p$=0.40, Hedges'g=0.10). To further assess whether maternal pitch or loudness differed, we performed t-tests with bootstrapping correction (n=10000) across participants for each metric. The results revealed that there was a significant difference across conditions for loudness ($p$<0.01) but not for pitch ($p$=0.1).

| | **Mean diff (Pos-Neg)** | **SD** | ***p*** | **t** | **df** | **Hedges'g** |
|---|---|---|---|---|---|---|
| **Loudness** | 60.766 | 38.38 | **0.001**** | 5.708 | 12 | 1.6093 |
| **Pitch** | 2.009 | 4.11 | 0.101 | 1.761 | 12 | 0.47014 |

**b) Inter-brain ANCOVA connectivity analysis**

Since loudness differed significantly across *Pos* and *Neg* conditions (whilst duration and pitch did not), we introduced loudness as a covariate in the inter-brain connectivity analyses for Strength, Divisibility and Inter-Brain Density (IBD). In the following, all calculations were corrected by bootstrapping (n=1000).

**i. Strength:**

| PDC | F (1,22) | *p* | $\eta^2$ | PLV | F(1,22) | *p* | $\eta^2$ |
|---|---|---|---|---|---|---|---|
| **Loudness** | 0.052 | 0.802 | 0.002 | **Loudness** | 0.208 | 0.565 | 0.009 |
| **Condition** | 2.819 | **0.019*** | 0.109 | **Condition** | 27.602 | **0.001**** | 0.545 |

For both PDC and PLV metrics, loudness did not emerge as a significant covariate in the inter-brain Strength ANCOVA analysis ($p>0.5$). Significant differences across conditions were still observed for both PDC and PLV metrics ($p<0.05$ and $p<0.01$ respectively) even after adding loudness as a covariate, replicating the results reported in the main text.

**ii. Divisibility:**

| PDC | F (1,23) | *p* | $\eta^2$ | PLV | F(1,23) | *p* | $\eta^2$ |
|---|---|---|---|---|---|---|---|
| **Loudness** | 0.368 | 0.509 | 0.016 | **Loudness** | 1.052 | 0.300 | 0.044 |
| **Condition** | 1.149 | 0.103 | 0.048 | **Condition** | 18.131 | **0.005**** | 0.441 |

For both PDC and PLV metrics, loudness did not emerge as a significant covariate in the inter-brain Divisibility ANCOVA analysis ($p>0.3$). After adding loudness as a covariate, there was still a significant difference across conditions for the PLV metric ($p<0.01$), in line with the results reported in the main text. However, for the PDC metric, there was now only a trend toward significance ($p=0.1$).

### ii. Inter-brain density (IBD):

| PDC | F (1,23) | *p* | $\eta^2$ | PLV | F(1,23) | *p* | $\eta^2$ |
|---|---|---|---|---|---|---|---|
| **Loudness** | 3.848 | 0.062 | 0.143 | **Loudness** | 3.052 | 0.094 | 0.117 |
| **Condition** | 0.273 | 0.610 | 0.012 | **Condition** | 5.687 | **0.005**** | 0.198 |

Finally, loudness was a marginally-significant covariate in the ANCOVA analysis of inter-brain density for both PLV and PDC ($p$<0.1). However, the addition of this covariate did not change the results as reported in the main text. Namely, there was still a significant difference between *Pos* and *Neg* conditions for the PLV metric ($p$<0.01), but no difference for the PDC metric ($p$>0.6), see Figure 8.
